# NusG is an intrinsic transcription termination factor that stimulates motility and coordinates global gene expression with NusA

**DOI:** 10.1101/2020.08.07.241513

**Authors:** Zachary F. Mandell, Reid T. Oshiro, Alexander V. Yakhnin, Mikhail Kashlev, Daniel B. Kearns, Paul Babitzke

## Abstract

NusA and NusG are transcription elongation factors that stimulate RNA polymerase pausing in *Bacillus subtilis*. While NusA was known to function as an intrinsic termination factor, the role of NusG in this process had not been explored. To examine the individual and combinatorial roles that NusA and NusG play in intrinsic termination, Term-seq was conducted in wild type, NusA depletion, Δ*nusG*, and NusA depletion Δ*nusG* strains. We determined that NusG functions as an intrinsic termination factor that works alone and cooperatively with NusA to facilitate termination at 88% of the 1,400 identified intrinsic terminators. The loss of both proteins leads to global misregulation of gene expression. Our results indicate that NusG stimulates a sequence-specific pause that assists in the completion of suboptimal terminator hairpins with weak terminal A-U and G-U base pairs at the bottom of the stem. Moreover, the loss of NusG results in flagella and swimming motility defects.

## Introduction

The transcription cycle can be subdivided into initiation, elongation, and termination. Regulation of initiation by a variety of DNA binding proteins is well established, while elongation is regulated by auxiliary transcription factors that interact with RNA polymerase (RNAP), including the elongation factors NusA and NusG (*Mondal et al., 2017*). Transcription termination demarcates the 3’ ends of transcription units, and misregulation of this process can result in spurious sense and/or antisense transcription (*Roberts, 2019; Mondal et al., 2016*). Termination in bacterial systems is known to proceed via two distinct mechanisms. One mechanism is Rho-dependent termination, which requires the activity of Rho, a hexameric ATP-dependent RNA translocase (*Roberts, 2019*). The other mechanism is intrinsic termination, which is generally assumed to not require the activity of additional protein factors and as such is also frequently referred to as factor-independent termination (*Roberts, 2019*). An intrinsic terminator is composed of a GC-rich hairpin followed immediately by a U-rich tract, both of which define the point of termination (POT) (*Roberts, 2019*). Completion of the terminator hairpin can induce transcript release via hybrid shearing and/or hyper-translocation depending on the transcriptomic context (*Roberts, 2019*).

NusA is a conserved bacterial transcription elongation factor that is essential for cellular viability in both *B. subtilis* and *Escherichia coli*. This protein factor binds to the flap-tip domain of the β subunit of RNAP via its N-terminal domain (NTD) (*Guo et al., 2018*). Once bound, NusA can directly interact with RNA using its S1, KH1 and KH2 domains (*Guo et al., 2018*). The binding of these domains to RNA elements allows NusA in combination with several other Nus proteins to serve as an antitermination factor when transcribing rRNA or bacteriophage λ sequences (*Nodwell et al., 1991; Vogel et al., 1997*). Also, the NTD of NusA can provide an additional set of positively charged residues outside of the RNA exit channel, extending this cavity while stabilizing the nucleation of RNA hairpins (*Guo et al., 2018*). Moreover, the binding of NusA results in an allosteric widening of the RNA exit channel of RNAP, which can then more readily accommodate RNA duplexes (*Guo et al., 2018*). The ability of NusA to promote the nucleation and formation of hairpins within the RNA exit channel allows this factor to serve as both a pausing factor and a termination factor, yet the relationship between the termination and antitermination activities of NusA is not well understood (*Guo et al., 2018*).

NusG, or SPT5 in archaeal and eukaryotic organisms, is the only universally conserved transcription factor. Bacterial NusG has two domains; the N-terminal NGN domain, which binds to the clamp helices of the β’ subunit of RNAP, and the KOW domain, which is connected to the NGN domain via a flexible linker and is free to interact with various regulatory partners (*Liu et al., 2017*). In *E. coli,* NusG is an anti-pausing factor and a core regulator of transcriptional polarity via the ability of its KOW-domain to interact with either Rho or ribosomal protein S10 (*Mooney et al., 2009; Tomar et al., 2013*). Also, the KOW domain of *E. coli* NusG can interact with the S1 domain of NusA when coordinated by NusE and λN during bacteriophage λ antitermination (*Krupp et al., 2019*). In contrast, *B. subtilis* NusG is a sequence-specific pausing factor due to the ability of its NGN domain to make direct contacts with the non-template DNA (ntDNA) strand within the transcription bubble (*Yakhnin et al., 2016*). This interaction results in a pause when the NGN domain encounters a stretch of T residues at critically conserved positions, as observed in 1,600 NusG-dependent pause sites genome-wide (*Yakhnin et al., 2020*). T residues in the ntDNA strand correspond to U resides in the nascent transcript. Although pausing is thought to be a fundamental prerequisite to termination, the role of NusG in intrinsic termination has not been investigated in *B. subtilis*.

While the current dogma posits that intrinsic termination generally does not require additional auxiliary protein factors, it was recently determined that NusA stimulates intrinsic termination on a global level in *B. subtilis*, with 232 intrinsic terminators classified as NusA-dependent (*Mondal et al., 2016)*. Our results show that *B. subtilis* NusG is also an intrinsic termination factor, and that NusA and NusG cooperatively stimulate intrinsic termination on an unexpectedly large scale, with only 12% of all identified intrinsic terminators continuing to terminate efficiently in the absence of these two proteins. Our results suggest a model in which NusG-dependent pausing plays a vital role in NusG-dependent termination, and that the absence of NusG results in the misregulation of global gene expression and altered cellular physiology and behavior.

## Results

### NusG and NusA cooperatively stimulate intrinsic termination *in vivo*

For this study, we used a *nusA*_dep_ strain in which NusA was solely generated exogenously from an IPTG-inducible promoter (*Mondal et al., 2016*). Thus, growth in the presence of IPTG results in wild type (WT) levels of NusA, whereas growth in the absence of IPTG results in near complete depletion of NusA within four cell generations as shown via Western blot (Supplementary Figure 1). By performing our studies with *nusA*_dep_ and *nusA*_dep_ Δ*nusG B. subtilis* strains ± IPTG we were able to mimic WT (*nusA*_dep_, +IPTG), NusA depletion (*nusA*_dep_, −IPTG), *nusG* deletion (*nusA*_dep_ Δ*nusG*, +IPTG), and NusA depletion *nusG* deletion (*nusA*_dep_ Δ*nusG*, −IPTG) conditions. To simplify the discussion, we will refer to these four conditions as WT, *nusA*_dep_, Δ*nusG*, and *nusA*_dep_ Δ*nusG* strains.

Term-seq is a bulk functional genomics assay that allows for the identification of all 3’ ends within a transcriptome via the ligation of a unique RNA oligonucleotide to the 3’ end of all transcripts isolated from a bacterial culture (*Mondal et al., 2016*). This ligation effectively preserves the authentic 3’ ends of all ligated transcripts, allowing for the computational identification of all authentic 3’ ends after sequencing (*Mondal et al., 2016*). To study the impact of NusA and NusG on intrinsic termination, we conducted Term-seq in the WT, *nusA*_dep_, Δ*nusG*, and *nusA*_dep_ Δ*nusG* strains. For each genomic region found to contain a stabilized transcript 3’ terminus, there were often multiple adjacent 3’ ends (*Mondal et al., 2016*). For our purposes, only the most abundant 3’ end within each region was included in the subsequent terminator analysis. Each 3’ end containing the core intrinsic terminator modules (RNA hairpin and U-rich tract) in the upstream sequence were categorized as potential intrinsic terminators, and a potential intrinsic terminator was confirmed to terminate *in vivo* only in cases where the termination efficiency (%T) at this nucleotide (nt) was ≥ 5 in the WT strain (see Materials and Methods). Using this system, we identified 4657 3’ ends in the WT strain (Supplementary Table 1), 1400 of which were categorized as intrinsic terminators (Supplementary Table 2). To benchmark the results of our assay, we compared the locations of all intrinsic terminators identified in this study, to all intrinsic terminators identified previously in WT *B. subtilis* grown in Minimal-ACH media (*Mondal et al., 2016*), and all intrinsic terminators identified by the *in silico* intrinsic terminator prediction tool TransTermHP applied to the *B. subtilis* genome (Supplementary Figure 2) (*Kingsford et al., 2007*). This analysis showed a high level of overlap between these datasets, with 937 terminators being conserved in all three datasets and 1329 terminators being shared between our dataset and at least one other dataset.

The %T was calculated for each of the 1400 intrinsic terminators in each strain, and a box plot was constructed to view the distribution of these data (Supplementary Table 2 and Figure 1A). NusG stimulated intrinsic termination to a similar extent as NusA, with a ~22% drop in median %T in the *nusA*_dep_ and Δ*nusG* strains when compared to the WT strain. Loss of both NusA and NusG in the *nusA*_dep_ Δ*nusG* strain resulted in a drastic termination defect, with the median %T falling 55%. Change in %T upon the loss of NusA and/or NusG (Δ%T) was calculated for all intrinsic terminators in each mutant strain (Supplementary Table 2, Column F). Based on our previously established categorization scheme, an intrinsic terminator was categorized as ‘dependent’ on NusA and/or NusG when the Δ%T ≥ 25, and ‘independent’ of NusA and/or NusG when 10 ≥ Δ%T ≥ −10 (*Mondal et al., 2016*). Surprisingly, only 12% of all intrinsic terminators were categorized as independent in the *nusA*_dep_ Δ*nusG* strain. To further assess the scope of the relationship between NusA and NusG on intrinsic termination, the overlap of intrinsic terminators categorized as dependent in each strain was organized into a Venn diagram and various intrinsic terminator subpopulations were identified (Figure 1B). Terminators that were classified as dependent in only the *nusA*_dep_ single mutant and *nusA*_dep_ Δ*nusG* double mutant strains were categorized as requiring NusA (Req A), while terminators that were classified as dependent in only the Δ*nusG* single mutant and *nusA*_dep_ Δ*nusG* double mutant strains were categorized as requiring NusG (Req G). Terminators that were categorized as dependent in all three strains were classified as requiring both NusA and NusG (Req A and G). A large number of terminators were only categorized as dependent in the double mutant strain, indicating they were able to terminate efficiently when either NusA or NusG was present in the cell, but not when both were absent. As such, this subpopulation was categorized as requiring either NusA or NusG (Req A or G). Intriguingly, 65% of all intrinsic terminators depicted in Figure 1B are present in the Req A and G or in the Req A or G subpopulations, clearly illustrating a large functional overlap between NusA and NusG on intrinsic termination.

**Figure 1.**
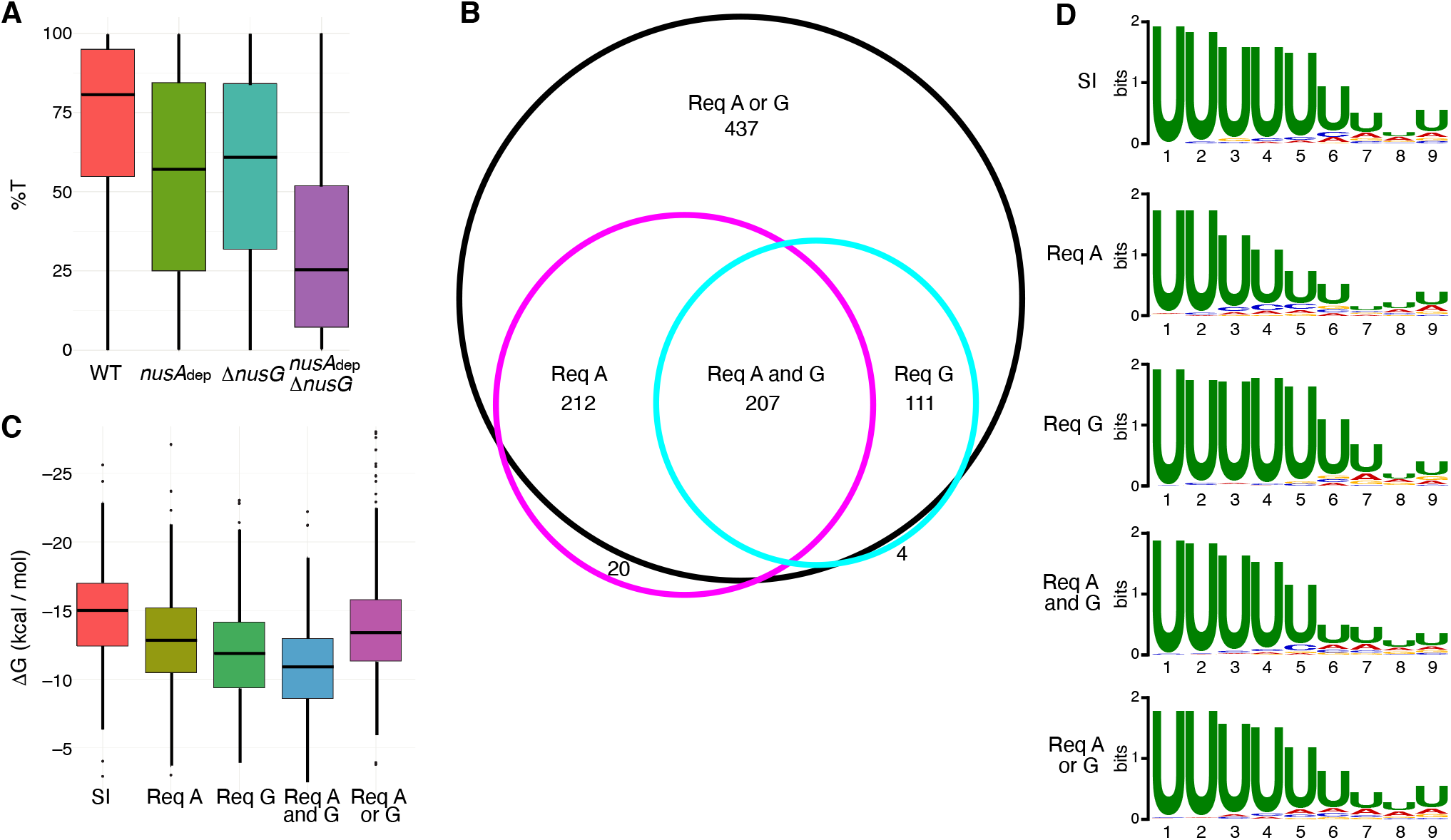
NusG is an intrinsic termination factor that works with NusA to stimulate suboptimal terminators. (**A**) Box plot showing the distribution of termination efficiency (%T) in WT, *nusA*dep, Δ*nusG*, and *nusA*dep Δ*nusG* strains. For this and all box plots, boundaries of the box designate the interquartile range (IQR), while upper/lower whiskers extend from the 75th/25th percentile to the largest/smallest value no further than 1.5*IQR in either direction. (**B**) Venn diagram showing the number and overlap of terminators that were classified as being dependent (Δ%T ≥ 25) on NusA (Req A) and/or NusG (Req G). The black circle contains all terminators that were classified as dependent in the *nusA*dep Δ*nusG* strain, the magenta circle contains all terminators that were classified as dependent in the *nusA*dep strain, and the cyan circle contains all terminators that were classified as dependent in the Δ*nusG* strain. Intrinsic terminator subpopulations that require NusA and/or NusG in any fashion to terminate efficiently are specified. (**C**) Box plot showing the distribution of predicted hairpin strength as reported in ΔG (kcal/mol) for all identified subpopulations including the strong and independent (SI) terminators. (**D**) Sequence logos of the U-rich tracts generated from the 9 nt window downstream of the predicted hairpins for all identified subpopulations including the strong and independent (SI) terminators.

The predicted hairpin stability for all intrinsic terminators within each identified subpopulation, including the subpopulation found to terminate strongly in the WT strain (%T ≥ 70) and be independent of both protein factors (strong and independent, SI), was calculated and organized into a box plot (Figure 1C). In parallel, sequence logos were generated from the 9 nt region immediately downstream of the predicted hairpin for terminators within each subpopulation (i.e. the predicted U-rich tract). Our terminator prediction system considered the hairpin to end before the first U residue and any contribution of A-U base pairing to terminator hairpin stability was not considered. It was found that terminators that require NusG in any fashion have weaker predicted hairpins than SI terminators, akin to Req A terminators, while terminators that require both NusA and NusG have the weakest hairpins. In addition, terminators that require NusG in any fashion exhibit a stronger enrichment of U residues immediately downstream of the predicted hairpin than either SI or Req A terminators (Figure 1D). This strong U enrichment is in line with the hypothesis that NusG stimulates intrinsic termination through its role in pausing (see below). Analyzing the distribution of predicted terminator hairpin stem lengths from each subpopulation shows that SI terminators have the longest hairpin stems with a median of 10 nt, while Req A and G terminators have the shortest hairpin stems with a median of 8 nt (Supplementary Figure 3A). This difference likely contributes to the observation that SI terminators have the strongest hairpins while Req A and G terminators have the weakest hairpins. A similar analysis examining terminator hairpin loop length showed no appreciable difference between any subpopulation (Supplementary Figure 3B).

### The NGN-domain of NusG promotes pausing at terminators with weak terminal base pairs

Our *in vivo* results indicated that NusG stimulates intrinsic termination cooperatively with NusA. To examine this phenomenon further while exploring a potential mechanism, we cloned six terminators found to require NusA and/or NusG *in vivo* for *in vitro* experimentation. We first examined the *yetJ* (Req A and G) (Figures 2A and 2B) and the *ktrD* (Req G) (Figures 2D and 2E) terminators. To maintain a logical consistency between our *in vivo* and *in vitro* data, each *in vivo* condition is labeled to indicate the elongation factors that were present in the cell. Single-round termination assays were conducted with these two terminators ± NusA and/or ± NusG. Experiments were also performed with the NusG NGN domain because this domain was found to be sufficient to recapitulate all features of NusG-dependent pausing (*Yakhnin et al., 2016*). NusG recapitulated the stimulatory effect on termination of the *yetJ* and *ktrD* terminators, as well as the cooperative effect between NusA and NusG that we observed *in vivo* (Figures 2C and 2F). Moreover, the highly similar termination patterns observed when using either full-length NusG or the NGN domain (± NusA) demonstrates that this phenomenon can be fully attributed to the NGN domain of NusG.

**Figure 2.**
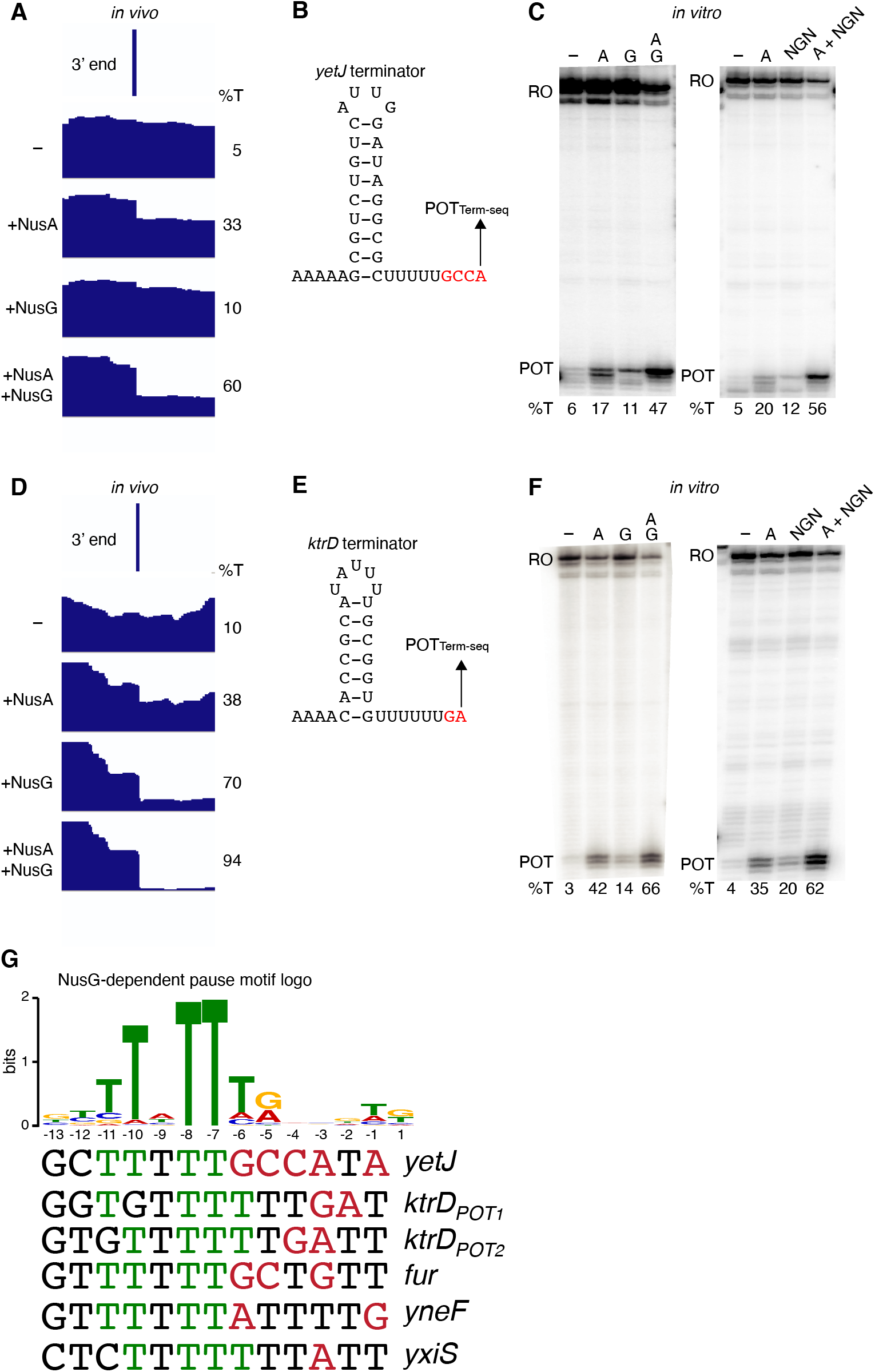
NusG stimulates intrinsic termination via its NGN domain. (**A**) IGV screenshot of a genomic window centered around the *yetJ* terminator. Top track is the 3’ end identified by Term-seq. Bottom tracks are the RNA-seq coverage data for the *nusA*dep Δ*nusG* (−), Δ*nusG* (+NusA), *nusA*dep (+NusG), and WT (+NusA +NusG) strains. %T in each strain is shown on the right of each track. Transcription proceeds from left to right. (**B**) *yetJ* terminator showing the point of termination identified by Term-seq *in vivo* (POT_Term-seq_). Disruptions in the U-rich tract are shown in red. The upstream A tract is also shown. (**C**) Single-round *in vitro* termination assay with the *yetJ* terminator. Experiments were performed in the absence (−) or presence of NusA (A), NusG (G), and/or NusG NGN domain as indicated. Positions of terminated (POT) and run-off (RO) transcripts are marked. %T is shown below each lane. (**D-F**) Identical to panels **A-C** except that it is the *ktrD* terminator. (**G**) NusG-dependent pause motif logo of the ntDNA strand is pictured on the top. ntDNA strand sequence upstream of the 3’ end identified by *in vitro* transcription for NusG-dependent terminators identified *in vivo*. Green nt are T residues that fit the NusG-dependent pause motif logo (TTNTTT). Red nt are non-U residues within the U-rich tract, which extends from positions −9 to −1.

Our results with the NGN domain suggested that NusG exerts its effect on intrinsic termination through its role in pausing. To test this hypothesis, single-round *in vitro* transcription time course (pausing) assays were conducted with the *ktrD* terminator ± NusG (Figures 3A and 3B). Results from this assay revealed 3 consecutive NusG-dependent transcription products that were 9, 10, and 11 nt downstream from the predicted terminator hairpin. Comparing the RNA species in the time course lanes with the 30 min termination lane shows that the product 9 nt from the predicted hairpin is a NusG-dependent termination site (POT_1_), the product 10 nt from the predicted hairpin is a NusG-dependent pause site and a NusG-dependent termination site (Pause_1_/POT_2_), and the product 11 nt from the predicted hairpin is a NusG-dependent pause site (Pause_2_).

**Figure 3.**
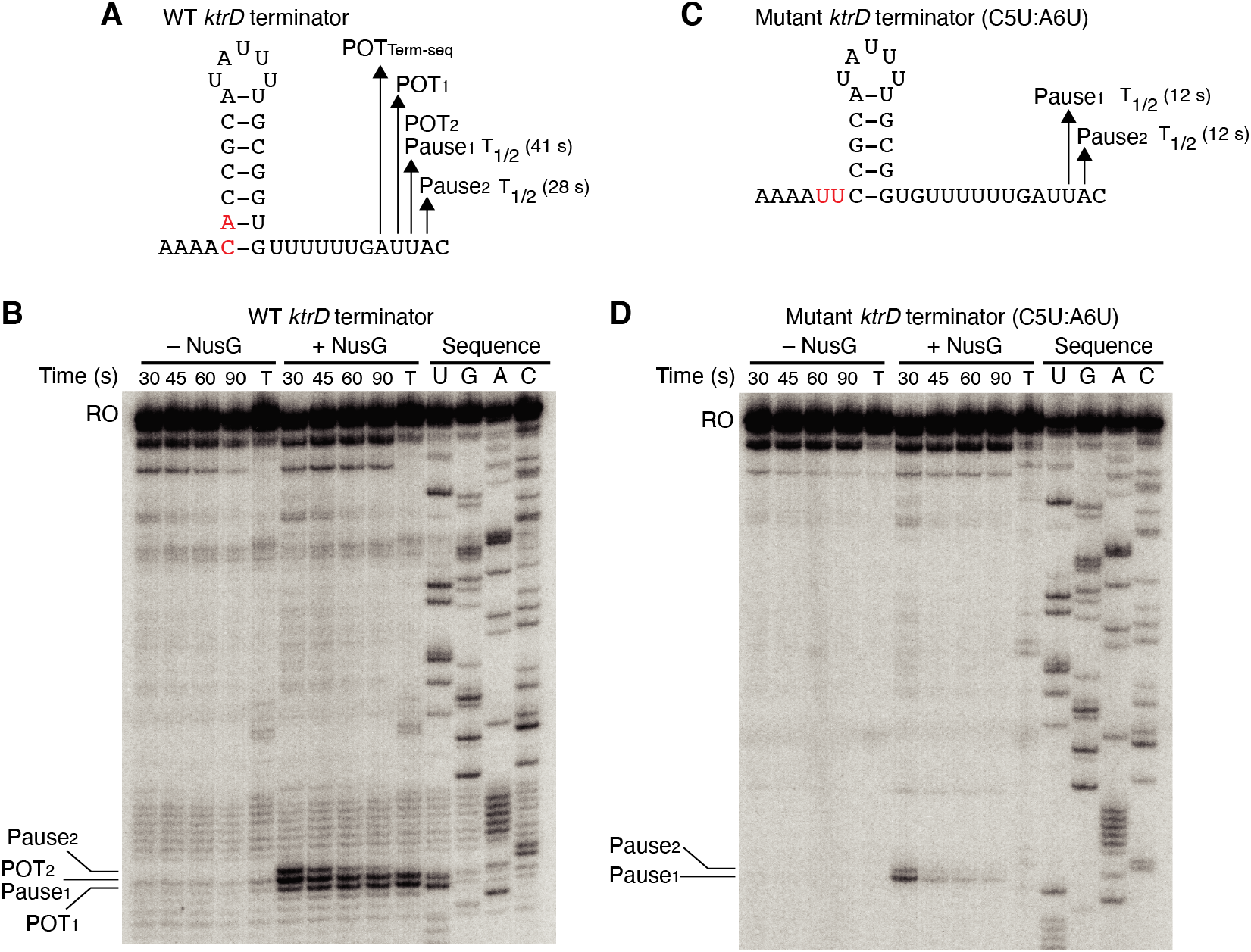
NusG stimulates termination through its role as a sequence-specific pause factor. (**A**) *ktrD* terminator showing the points of termination (POT1 and POT2) and pause sites (Pause1 and Pause2) identified *in vitro*. The POT identified *in vivo* by Term-seq is also specified (POT_Term-seq_). The upstream A tract is also shown. The half-life (T1/2) of each pause based on data within panel **B** are specified to the right of each pause in parentheses. (**B**) Single round *in vitro* pausing and termination assay using the WT *ktrD* terminator ± NusG. Time points of elongation are indicated above each lane. T, 30 min termination assay. RNA sequencing lanes (U, G, A, C) are labeled. Positions of NusG-dependent pause bands, termination sites, and run-off (RO) transcripts are marked. (**C, D**) Identical to **A-B** except that it is the C5U:A6U mutant *ktrD* terminator. The WT (**A**) and mutated (**C**) residues are highlighted in red.

The hairpin to 3’ end distance for intrinsic terminators is 7-9 nt, whereas this distance is 11-12 nt for pause sites (*Mondal et al., 2016; Yakhnin et al., 2020; Ray-Soni et al., 2016*). Our results with the *ktrD* terminator indicate that NusG can elicit a pause at two consecutive positions, but only one of these positions contains the requisite elements to induce transcript release. This distinction likely depends on the hairpin to 3’ end distance achieved at each position before readthrough. To test this possibility, we mutated the *ktrD* terminator by substituting C5 and A6 with U residues to reduce the length of the hairpin and to increase the hairpin to 3’ end distance by 2 nt while leaving the NusG-dependent pause motif intact (Figure 3C). These mutations altered the transcription profile of this template such that pausing still occurred at positions identical to the positions of Pause_1_ and Pause_2_ on the WT template (Figure 3D). These results indicate that the NusG-dependent pause motif upstream of the 3’ end is sufficient to elicit a pause and that the position of this pause is set by the motif. Calculating the half-life of both pauses on each template revealed that the duration of the pauses on the mutant template was shorter than the pauses on the WT template, implying that a NusG-dependent pause can be stabilized to different degrees by hairpins of different lengths and strengths. Interestingly, a fraction of RNAP at Pause_1_ on the mutant template remained paused until at least 90 s, suggesting that NusG-dependent pausing at this position is highly stable. Termination no longer occurred on the mutant template, demonstrating that we successfully converted a NusG-dependent terminator into a NusG-dependent pause site by extending the hairpin to 3’ end distance by 2 nt.

Termination occurring 7-9 nt downstream of the hairpin is a biophysical constraint set by the length of the RNA-DNA hybrid (*Ray-Soni et al., 2016*). The presence of terminated RNA species 10 nt downstream of the predicted *ktrD* terminator hairpin implies that this hairpin in reality extends further via A-U base pairing. Terminators with hairpin stems that contain multiple consecutive terminal A-U base pairs are highly atypical, yet a similar phenomenon was observed for the *yetJ* (Req A and G), *fur* (Req A and G), *yneF* (Req A and G), and *yxiS* (Req G) terminators (Figure 4 and Supplementary Figure 4). Notably, in all cases NusG effectively stimulated termination at a POT that utilized hairpins with 2 to 4 consecutive terminal A-U base pairs. Surprisingly, we also found that NusG stimulated the *fur* terminator to terminate at a position 9 nt downstream of a hairpin that utilized a terminal G-U base pair (Figure 4). This combination of terminator features has never before been reported within the same terminator.

**Figure 4.**
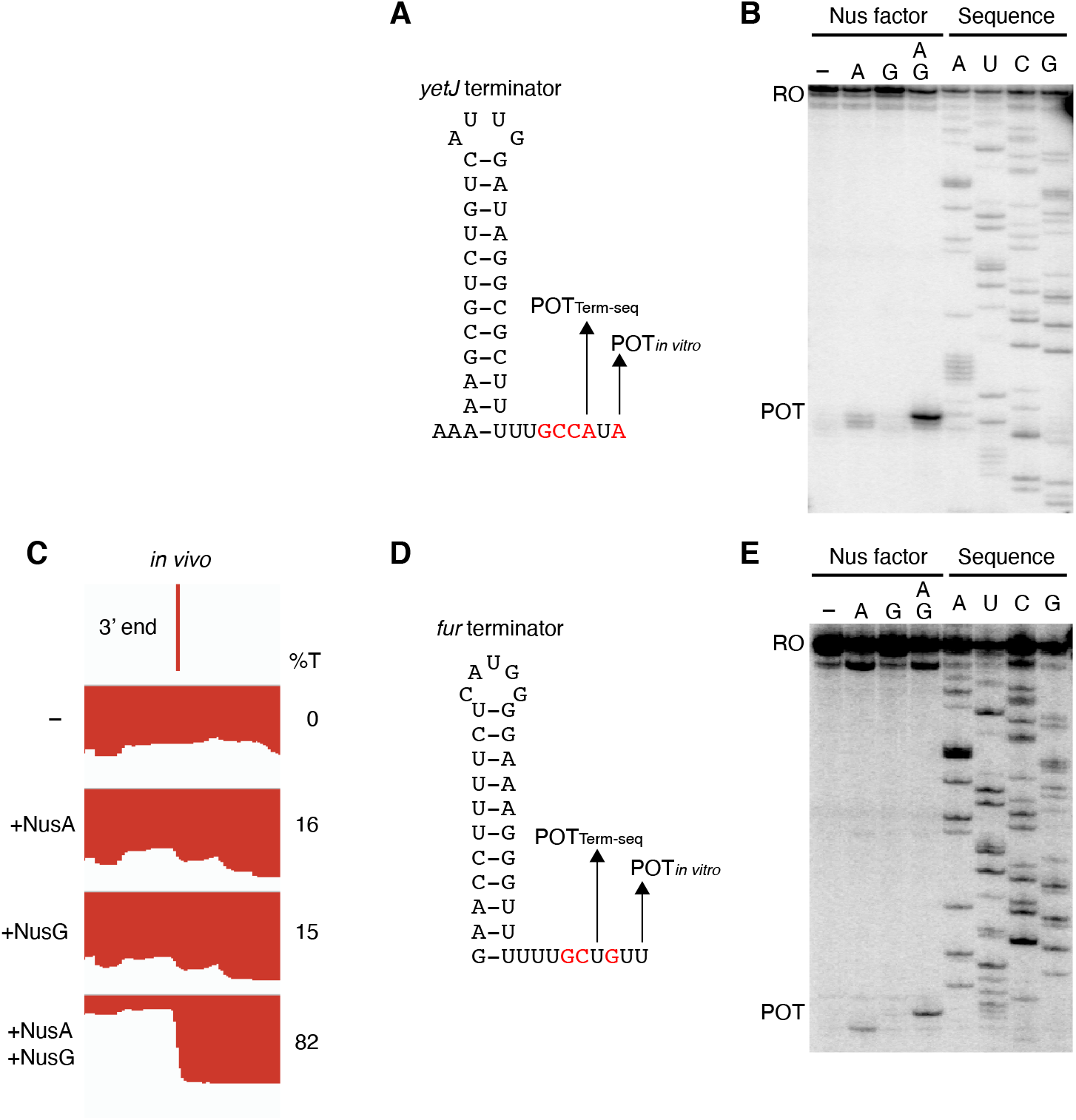
NusG stimulates terminators with particularly weak terminal base pairings. (**A**) *yetJ* terminator showing the point of termination identified *in vivo* by Term-seq (POT_Term-seq_) and by *in vitro* transcription in the +A+G condition (POT_*in vitro*_). Disruptions in the U-rich tract are shown in red. The upstream A tract is also shown. An IGV screenshot of this terminator is shown in Figure 2A. (**B**) Single-round *in vitro* termination assay with the *yetJ* terminator. Experiments were performed in the absence (−) or presence of NusA (A) and/or NusG (G) as indicated. Positions of terminated (POT) and run-off (RO) transcripts are marked. RNA sequencing lanes (A, U, C, G) are labeled. (**C**) IGV screenshot of a genomic window centered around the *fur* terminator. Top track is the 3’ end identified by Term-seq. Bottom tracks are the RNA-seq coverage data for the *nusA*dep Δ*nusG* (−), Δ*nusG* (+NusA), *nusA*dep (+NusG), and WT (+NusA +NusG) strains. %T in each strain is shown on the right of each track. Transcription proceeds from right to left. (**D, E**) Identical to panels **A, B** except that it is the *fur* terminator. (**D**) Note that the terminal 3 base pairs contain the A tract and one G residue.

While the sequence logos generated from our *in vivo* terminator hairpin predictions shows an enrichment of U residues immediately downstream of the predicted hairpin for terminators that depend on NusG (Figure 1D), our *in vitro* 3’ end mapping suggests that the upstream portion of this U-rich tract is actually present in the base of the terminator hairpin. Comparison of the NusG-dependent pause motif with the terminator sequences confirmed *in vitro* shows that each terminator contains U residues at positions that correspond to the T residues in the ntDNA strand that are most critical for NusG-dependent pausing (Figure 2G, green residues) (*Yakhnin et al., 2020*). Moreover, the updated U-rich tract sequences shows that NusG-dependent terminators frequently contain distal U-rich tract interruptions, akin to NusA-dependent terminators (Figure 2G, red residues) (*Mondal et al., 2016*).

### NusG and NusA cooperatively coordinate global gene expression

Terminator read-through can impact gene expression by increasing transcription of downstream genes oriented in the same direction, destabilizing transcripts from downstream convergent transcription units, and/or changing the expression of global regulators (*Mondal et al., 2016*). To examine the effect of NusG on gene expression, a differential expression analysis was conducted comparing expression data from each mutant strain to expression data from the WT strain (Supplementary Table 3). Volcano plots were constructed based on the results of this analysis, and affected genes were determined using false discovery rate (FDR) cutoffs of 0.005 and fold-change cutoffs of 4. This approach revealed that NusG is involved in regulating gene expression on a global scale, with 106 transcripts increasing in expression and 37 transcripts decreasing in expression in the Δ*nusG* strain (Figure 5A). NusA had a larger effect on gene expression with 322 transcripts increasing in expression and 94 transcripts decreasing in expression in the *nusA*_dep_ strain (Figure 5B). The cooperative relationship between those two proteins on intrinsic termination extended to gene expression, with 28% of all transcripts expressed in the WT strain being misregulated in the *nusA*_dep_ Δ*nusG* strain (Figures 5C and 5D).

**Figure 5.**
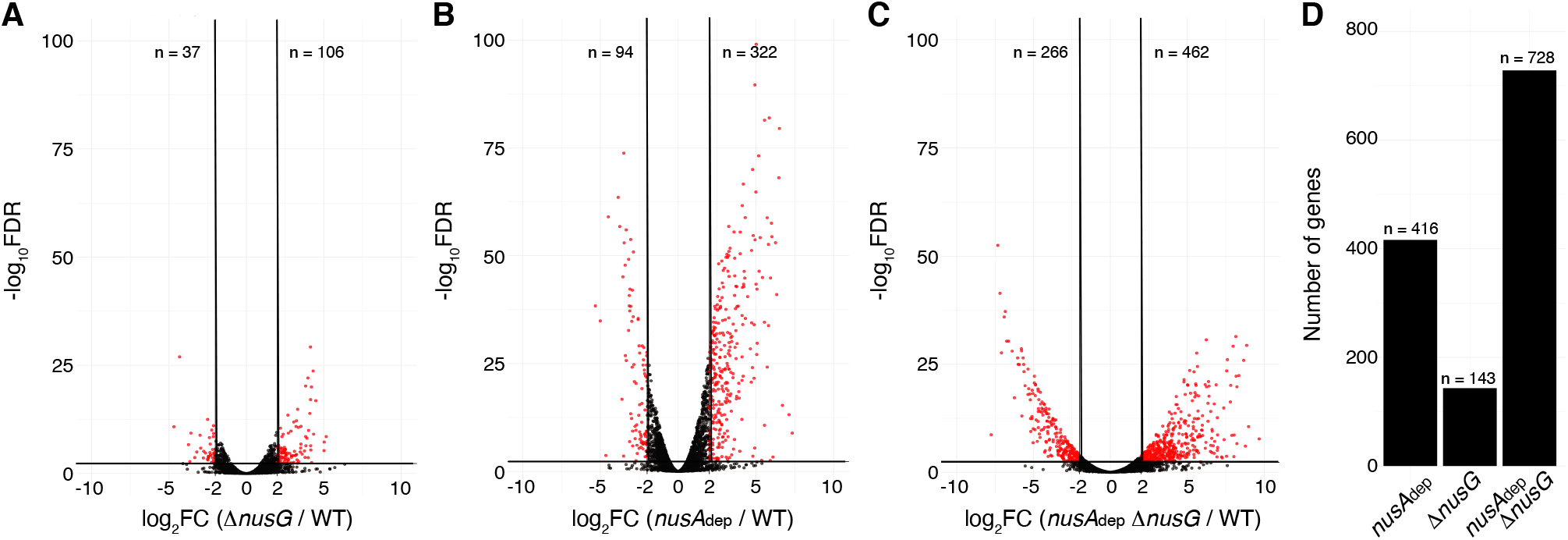
NusG coordinates global gene expression with NusA. (**A**) Volcano plot derived from differential expression analysis comparing steady-state gene expression levels in the WT and *ΔnusG* strains. Cutoffs are Log2 fold-change (Log2FC) of 2 and a −Log10 of the False Discovery Rate (-Log10FDR) of 2.3. Number (n) of genes downregulated and upregulated are specified. (**B**) Identical to panel **A** except comparing the WT and *nusA*dep strains. (**C**) Identical to panel **A** except comparing the WT and *nusA*dep Δ*nusG* strains. (**D**)Total number of differentially expressed genes in the *nusA*dep, Δ*nusG*, and *nusA*dep Δ*nusG* strains. 2,604 transcripts in which the transcripts per million (TPM) > 10 in the WT strain were used in this analysis (Supplementary Table 4, column D).

### NusG plays a critical role in regulating swimming motility

The transcripts per million (TPM) of genes within the motility, sporulation, and competence regulons were calculated for each strain, and the mutant TPM values were compared to their WT counterparts (Supplementary Table 4). Loss of NusG resulted in a median 3-fold decrease in expression of genes within the motility regulon, while loss of both NusA and NusG resulted in a median 8-fold decrease in expression of these genes (Figure 6A). The 27 kb *fla*/*che* operon is composed of 32 genes involved in flagella biosynthesis and motility (*Cozy et al., 2010; Márquez-Magaña et al., 1994; Albertini et al., 1991*). Transcription of the majority of genes within the motility regulon, including the *fla*/*che* operon, is initiated by RNAP directed by the alternative sigma factor σ^D^, which is encoded by the penultimate gene within the *fla*/*che* transcript, *sigD* (*Helmann et al., 1988; Márquez et al., 1990; Serizawa et al., 2004; Kearns et al., 2005*). To assess the expression trends of the *fla*/*che* operon, expression of the 2^nd^ gene of the cluster (*flgC*) and *sigD* were calculated, showing a 1.7-fold and 3-fold decrease in the expression of *flgC* and *sigD* in the Δ*nusG* strain and a 2.1-fold and 10-fold decrease in the expression of *flgC* and *sigD* in the in the *nusA*_dep_ Δ*nusG* strain (Figures 6B and 6C). As such, the ratio of *flgC* to *sigD* expression increased from 1.3-fold in the WT strain, to 2.1-fold in the Δ*nusG* strain and 6.2-fold in the *nusA*_dep_ Δ*nusG* strain (Figures 6B and 6C). Higher expression of the 5’ portion of a transcript compared to the 3’ portion of a transcript is sometimes caused by the activity of 3’ to 5’ exoribonucleases such as PNPase and RNase R (*Liu et al., 2014; Baumgardt et al., 2018; Bechhofer et al., 2019*). While there were moderate increases in the expression of the genes encoding for PNPase (*pnpA*) and RNase R (*rnr*) in the absence of NusG (Supplementary Figure 5A), the changes in expression of *pnpA* were not considered statistically significant during the differential expression analysis (Supplementary Table 3, Column G) and the expression of *rnr* remained relatively low in all conditions (Supplementary Figure 5A). Thus, the observed increase in the ratio of *flgC* to *sigD* expression cannot be fully explained by an increase in known mediators of RNA decay, and is likely due to a defect in transcript completion, which has been posited to impact the expression of *sigD* compared to the 5’ portion of the *fla*/*che* operon (*Cozy et al., 2010*).

**Figure 6.**
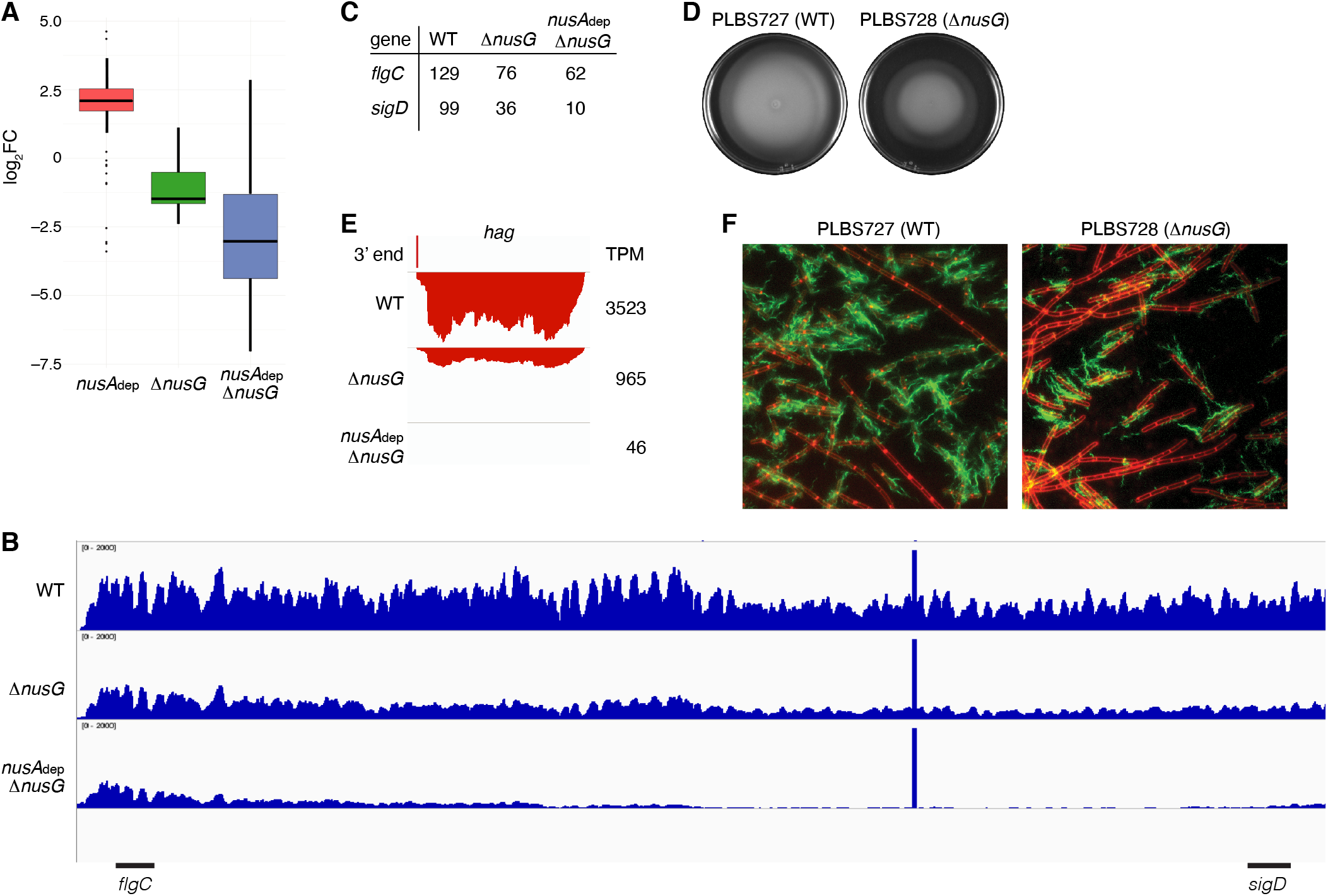
NusG is a motility factor in *B. subtilis*. (**A**) Box plot showing the effect of NusA and/or NusG for all transcripts within the motility regulon; log2FC (mutant:WT). (**B**) IGV screenshot of the *fla/che* operon. Each track is the RNA-seq coverage data for the WT, Δ*nusG*, and *nusA*dep Δ*nusG* strains. Locations of the *flgC* and *sigD* genes are specified below the screenshot. (**C**) TPM values calculated for the *flgC* and *sigD* genes in the WT, Δ*nusG*, and *nusA*dep Δ*nusG* strains. (**D**) Swimming motility assay for PLBS727 (WT) and PLBS728 (Δ*nusG*). (**E**) IGV screenshot of the *hag* transcript. Top track is the 3’ end identified by Term-seq. Bottom tracks are the RNA-seq coverage data for the WT, Δ*nusG*, and *nusA*dep Δ*nusG* strains. Transcription proceeds from right to left. TPM values were calculated for the *hag* transcript in each strain and are specified to the right of each track. (**F**) Fluorescence microscopy performed on PLBS727 (WT) and PLBS728 (Δ*nusG*) strains. Membrane is stained with FM4-64 (false colored red) and flagella are stained with Alexa Fluor 488 C5 maleimide (false colored green).

To explore how the changed expression of the *fla*/*che* transcript effected motility, a swimming motility assay was conducted on WT, *nusA*_dep_, Δ*nusG*, and *nusA*_dep_ Δ*nusG* strains (Figure 6D and Supplementary Figure 5B). To eliminate complications in the interpretation of the assay caused by the differing growth rates of the NusA depleted strains, the assay was also conducted on strains PLBS727 (WT *nusG* and WT *nusA*) and PLBS728 (Δ*nusG* and WT *nusA*) (Figure 6D). Loss of NusG in our Δ*nusG* strain and in PLBS728 resulted in an impaired swimming motility phenotype. This impaired swimming motility phenotype may have been further exacerbated by depletion of NusA, although the reduced growth rate of the NusA depleted strains complicated interpretation (Supplementary Figure 5B).

The *hag* transcript, encoding the flagellin protein in *B. subtilis,* is one of the most abundant transcripts in the cell, and its expression is entirely dependent on σ^D^-directed RNAP (*Mondal et al., 2016; LaVallie et al., 1989; Mirel et al., 1989*). Consistent with a cascading reduction in expression of the σ^D^ regulon due to the reduction of *sigD* expression, expression of *hag* mRNA was reduced 4-fold in the Δ*nusG* strain and by an astonishing 75-fold in the *nusA*_dep_ Δ*nusG* strain (Figure 6E). To monitor how the reduction of *hag* expression impacts Hag production, we engineered a Hag variant with a surface exposed cysteine residue to be expressed ectopically from its native promoter. This Hag variant could then be fluorescently labeled via a cysteine-reactive fluorescent maleimide dye (*Blair et al., 2008*). Introducing this allele into the same six strains that were used for the swimming motility assay allowed us to determine whether the absence of NusA and/or NusG effected flagella synthesis levels (Figure 6F and Supplementary Figure 5C). Through this experiment, we observed both a decrease in the frequency of cells that produced flagella and a reduced number of flagella per cell. We conclude that the loss of NusG results in a motility defect that was correlated with a reduction in the expression of the *fla/che* operon, *hag*, and the majority of the σ^D^ regulon.

## Discussion

In this work we conducted Term-seq in WT, *nusA*_dep_, Δ*nusG*, and *nusA*_dep_ Δ*nusG* strains of *B. subtilis.* By locating all intrinsic terminators in the WT condition, we were able to singly and combinatorially quantify the effect of NusA and NusG on intrinsic termination. Although NusG was not recognized as an intrinsic termination factor prior to these studies, we found that NusG stimulates termination at intrinsic terminators with suboptimal hairpins and strong NusG-dependent pause signals upstream of the 3’ end. We also found that NusG works in a cooperative fashion with NusA to regulate both intrinsic termination and global gene expression. This cooperativity during termination can be fully recapitulated by the NGN domain of NusG *in vitro*. While the AR2 domain of NusA and the NGN domain of NusG were found to physically interact in *E. coli* during elongation, this interaction is unlikely to be relevant to the regulation of elongation in *B. subtilis* due to the absence of the AR2 domain in *B. subtilis* NusA (*Strauß et al., 2016*). Thus, *B. subtilis* NusA and NusG likely work together without physical interaction to cooperatively stimulate termination *in vivo*. Only 12% of all intrinsic terminators function independently of both NusA and NusG. Thus, our results establish that intrinsic, or factor-independent termination is primarily a factor-mediated process in *B. subtilis.* These findings are in stark contrast to the general dogma that intrinsic terminators are factor-independent and thus contain all elements required for transcript release within the nascent RNA sequence, RNA-DNA hybrid, and downstream DNA sequence. Thus, a new nomenclature system that divides intrinsic terminators based on their factor-dependency profiles is warranted. We suggest NusA-dependent terminators, NusG-dependent terminators, NusA-NusG-dependent terminators, and factor-independent terminators.

*In vitro* experimentation was conducted on 6 terminators that were found to be stimulated by NusG *in vivo*, and it was generally found that NusA is the more potent termination factor *in vitro* (Figures 2 and 4, Supplementary Figures 4 and 6). Interestingly, these templates included terminators that were found to be more reliant on NusG than NusA *in vivo.* This discrepancy could be explained by the inability of the *in vitro* assay to fully mimic *in vivo* conditions and/or the participation of additional protein factors *in vivo*. It was recently found that RNAP pausing at intrinsic terminators leads to RNAP collision at convergent transcription units and that these collisions can result in transcript release within *E. coli* (*Ju et al., 2019*). Knowing that pausing is involved in NusG-dependent intrinsic termination, we tested transcription *in vitro* on terminators that were both identified to be NusG-dependent and at which convergent transcription occurs *in vivo*. However, convergent transcription had no impact on the effect of NusG on termination *in vitro* (Supplementary Figure 7).

We found that NusG stimulated termination at a position further downstream in three templates tested *in vitro* (Figures 2C, 4B, and 4E). This finding can be explained by the fact that NusG shifts RNAP to the post-translocation register (*Yakhnin et al., 2020*) and may thus function as a processivity factor until encountering the consensus pause motif at these terminators. NusA has been reported to stimulate termination at a more upstream position (*Yakhnin et al., 2002*). This phenomenon was only observed *in vitro* in reactions where NusG was absent, implying that the shifting of RNAP to the post-translocation register is a central feature of elongation complexes containing NusG. Moreover, NusA and NusG together stimulated termination *in vitro* at a position 1 to 3 nt further downstream than the 3’ end identified by Term-seq *in vivo* on all templates tested (Figures 3B, 4B, 4E, and Supplementary Figures 4C and 4F). The discrepancy between the *in vivo* POT(s) and the *in vitro* POT(s) can perhaps be attributed to exoribonuclease trimming of the terminated transcripts *in vivo*, which has shown to be impeded by stem loops ≥ 7 bp in length (*Spickler et al., 2000*).

Our analysis indicates that NusG-dependent pausing is an essential component of certain intrinsic terminators. Furthermore, the feature that distinguishes a NusG-dependent terminator from a hairpin-stabilized NusG-dependent pause is whether the hairpin can extend to within 7-9 nt of the RNA 3’ end. To extend the time frame allowed for potential terminator hairpin completion, NusG elicits a sequence-dependent pause when RNAP incorporates U residues into the nascent transcript at positions −12, −11, −8, −7, and −6 relative to the 3’ end at position −1 (*Yakhnin et al., 2016; Yakhnin et al., 2020*). These U residues correspond to a TTNTTT consensus pause motif in the ntDNA strand of the paused transcription bubble (Figure 2G). Interaction of the NGN domain with these T residues is required for NusG-dependent pausing (*Yakhnin et al., 2016; Yakhnin et al., 2020*). We found that NusG is able to elicit a pause at intrinsic terminators, sometimes at several consecutive positions. Theoretically, NusG should be able to stimulate termination at any intrinsic terminator containing U residues at the −8, −7, and −6 positions. However, the only terminators that required NusG were those that have weak hairpins and U residues that extend upstream to the −11 and/or −12 positions and are thus the bottom-most base pairs of the hairpin stem (Figures 2G, 3, 4, and Supplementary Figure 4).

Terminator hairpins with weak terminal base pairs are highly atypical due to the various structural elements of RNAP that stabilize the elongation complex, including the interactions between the Sw3 pocket and the lid of RNAP with the −10 and −9 positions of the RNA transcript, respectively (*Ray-Soni et al., 2016*). Displacement of these interactions requires major energetic expenditures, which may explain why certain bacterial species evolved a strong G-C preference at the base of the terminator hairpin to drive completion of its formation (*Ray-Soni et al., 2016; Peters et al., 2011*). However, we found that the potential for additional A-U or G-U (A/G-U) base pairing was common in *B. subtilis*, with ~75% of all intrinsic terminators included in Figure 1b containing the potential for an A/G-U hairpin extension ≥ 2 nt in length (Supplementary Figure 8). The observation that NusG stimulates termination at a POT further downstream, while NusA stimulates termination at a POT further upstream implies that these A/G-U base pairs may only be bonafide members of the hairpin for terminators that depend on NusG.

The precise mechanism of transcript release at Nus-dependent terminators is not currently understood. One potential mechanism is that NusA and/or NusG simply assists with the formation of suboptimal terminator hairpins, the completion of which drives transcript release in all cases where the U-rich tract is at or above an optimality threshold. Additionally, the ability of NusG to shift RNAP into the post-translocation register may assist in the hyper-translocation and deactivation of RNAP at intrinsic terminators with weak terminal base-pairs and distal U-tract interruptions, while NusA assists in the formation of suboptimal hairpins and/or pausing at suboptimal U-rich tracts. The observation that NusA stimulates termination at a more upstream position provides evidence for a model in which terminators that depend on the formation of terminal A/G-U base-pairs require NusG, while terminators that have G-C or C-G terminal base-pairs may not be able to induce a sufficiently stable NusG-dependent pause, and therefore will depend solely on NusA.

Loss of NusG results in an impaired motility phenotype that can be traced back to a changed pattern in expression of the *fla*/*che* operon. A decrease in the expression of the 3’ portion of the *fla*/*che* operon compared to the 5’ portion was previously reported in *B. subtilis,* and this phenomenon may be caused by a stochastic release of RNAP during the transcription of this long operon (*Cozy et al., 2010*). The possibility that NusG functions as a processivity factor in *B. subtilis* may explain the potential defect in transcript completion upon the loss of NusG. Moreover, the fact that NusA depletion alone resulted in an increase in expression across the *fla*/*che* operon is suggestive of a generally destabilizing role of NusA on transcription (Supplementary Figure 9). The increase in the expression of the motility regulon in the NusA_dep_ strain can likely be explained by an increase in the expression of *sigD* (Figure 6A, Supplementary Figure 9) (*Estacio et al., 1998; West et al., 2000*). These results hint that modulating the activity of NusG and/or NusA governs the frequency of completion of the *fla/che* operon transcript and serves as a method for *B. subtilis* to regulate the switch between motile and sessile states.

*B. subtilis* NusG contacts the ntDNA strand to elicit a pause via an NT dipeptide located within the NGN domain (*Yakhnin et al., 2016*). A phylogenetic analysis focusing on bacterial species that contain a *B. subtilis*-like NusG homolog with an NT or HT dipeptide shows that the ability of NusG to contact the ntDNA strand may be present in a large number of Gram-positive and Gram-negative phyla (Supplementary Figure 10) (*Yakhnin et al., 2020*). Moreover, mycobacterial NusG was found to contain an NT dipeptide at this position and thus the discovery that mycobacterial NusG stimulates intrinsic termination at suboptimal terminators *in vitro* (*Czyz et al., 2014*) suggests that the NusG-dependent termination mechanism may be conserved as well.

## Methods

### *B. subtilis* strains

All strains used in this study are listed in Supplementary Table 6. Strains PLBS727 and PLBS728 are WT and Δ*nusG* strains, respectively. PLBS730 is a NusA depletion strain. This strain contains an IPTG-inducible *nusA* allele and *E. coli lacI* integrated into the chromosomal *amyE* gene (*Yakhnin et al., 2020*). PLBS731 is identical to PLBS730 except that it also contains Δ*nusG* (*Yakhnin et al., 2020*). NusA production was maintained in these two strains by culturing cells in the presence of 0.2 mM IPTG. PLBS730 and PLBS731 grown + 0.2 mM IPTG were considered to be WT and Δ*nusG,* respectively. These strains grown in the absence of IPTG were considered as *nusA*_dep_ and *nusA*_dep_ Δ*nusG,* respectively. The *lacA::Phag-hag^T209C^ tet* construct was generated by digesting *Phag-hag*^*T209C*^ fragment from plasmid pNE4 using BamHI and SphI (*Konkol et al., 2013*). The digested fragment was ligated into the BamHI and SphI sites of pNC018 (*lacA::tet*) to generate plasmid pKB141 (*Konkol et al., 2013*). Plasmid pKB141 was introduced into strain DS2569 by natural competence to generate strain DS6331, and further introduced into appropriate strain backgrounds by SPP1-mediated transduction (*Konkol et al., 2013*). All bacterial strains and plasmids are available from the corresponding author.

### *B. subtilis* growth and library generation

Each strain was streaked onto LB plates containing 20 μg/ml chloramphenicol and 0.2 mM IPTG. Single colonies of each strain were grown at 37°C as overnight standing cultures in 5 mL of LB media supplemented with 0.4 mM IPTG and 20 μg/ml chloramphenicol. The next day, 1 mL of cells were collected and washed twice with LB media. For strains to be depleted of NusA, a 30-fold dilution was made into 25 mL of LB supplemented with 20 μg/ml chloramphenicol (no IPTG). For strains in which NusA expression was maintained, a 75-fold dilution was made into 25 mL of LB media supplemented with 20 μg/ml chloramphenicol and 0.2 mM IPTG. All cultures were grown shaking at 37°C and both total RNA and total protein were extracted during mid-exponential phase. Barcoded Illumina libraries were generated from oligo-ligated transcripts as described previously (*Mondal et al., 2016*). Total RNA was CIP treated and rRNA was depleted from each sample using ribo-zero rRNA depletion kits (Illumina). The remaining transcripts were then ligated to a unique 2’, 3’ dideoxy RNA oligonucleotide (IDT) that was phosphorylated on the 5’ end. TruSeq standard mRNA libraries were generated from these samples. Equal amounts of each library were pooled and 150 nt single-read sequencing was performed with an Illumina NextSeq 500 in High Output mode.

### Western blot

NusA depletion was confirmed via western blot for all replicates (Supplementary Figure 1). Protein samples (3 μg) were fractionated in a 10% SDS gel and transferred to a 0.2 μm PVDF membrane. Purified his-tagged NusA, σ^A^, and cell lysates were probed with rabbit anti-NusA or anti-σ^A^ antibodies (1:5,000 dilution), and developed using enhanced chemiluminescence following incubation with HRP-conjugated goat anti-rabbit antibody (GenScript). Two images were taken of each membrane, one after probing for NusA and a second after probing for σ^A^.

### Data processing, analysis, and identification of 3’ ends

Illumina sequencing generated 141,842,192 reads across 8 samples (WT, *nusA*_dep_, Δ*nusG*, and *nusA*_dep_ Δ*nusG*). All reads were processed to comprehensively yield all 3’ ends as described previously with modifications (*Mondal et al., 2016*). After demultiplexing, Illumina adapters were trimmed with Trimmomatic, resulting in a traditional RNA-seq dataset (*Bolger et al., 2014*). Cutadapt was then used to extract all reads that were found to contain the unique RNA oligonucleotide used during library generation, resulting in a Term-seq dataset (*Marcel, 2011*), which was mapped to the *B. subtilis* 168 genome (NC_000964.3) via bwa-mem in single-end mode (*Li, 2013*). Bam files for each pair of replicates were merged, the contents of each resulting bam file was split by strand using samtools, and coverage files were generated for each strand-specific bam file using bedtools (*Li et al., 2009; Quinlan et al., 2010*). A series of custom python scripts were used to comprehensively identify all 3’ ends by calculating the coverage variation (C_V_) at each nt of all strand-specific coverage files, and identifying the local maxima across these C_V_ landscapes as described previously (*Mondal et al., 2016*). The C_V_ magnitude at a 3’ end is tightly correlated with 3’ abundance, which is a function of transcript abundance, 3’ end stability, and termination efficiency (%T) in cases where the 3’ end is a result of termination (*Mondal et al., 2016*). To limit 3’ ends that could be attributed to noise, a C_V_ threshold was set at ≥ 10 for this study. All strand-specific 3’ ends were then merged by biological condition, thereby generating the final 3’ end bedgraph files that contained the genomic location, the C_V_, and the strand information of each identified 3’ end (included with additional information in Supplementary Table 1). RNA-seq coverage files were generated by aligning the merged RNA-seq datasets to the *B. subtilis* genome using bwa-mem in single-end mode, splitting the resultant bam files by strand using samtools, using bedtools to calculate the per-nucleotide RNA-seq coverage, which were then merged by biological condition (*Li, 2013; Li et al., 2009; Quinlan et al., 2010*).

For construction of the phylogenetic tree, 10,000 NusG-homologs were identified by querying the NusG recognition region (DDSWXXVR<u>**XX**</u>PXVXGFXG) using BLASTp, where X indicates any amino acid and the underlined region is the dipeptide by which *B. subtilis* NusG uses to contact the ntDNA strand (*Yakhnin et al., 2016; Altschul et al., 1990*). From these 10,000 homologs, 776 representative bacterial genera were identified, 617 of which were found to contain a *B. subtilis*-like dipeptide within the underlined region (NT or HT), and these species were chosen to construct a 16S rRNA-based phylogeny using the NCBI common taxonomy tree webtool (*Sayers et al., 2009*). Tree annotation and display were created with the interactive tree of life web platform (iTOL) (*Letunic et al., 2016*). All sequence logos used in this study were generated using the MEME suite (*Bailey et al., 1994*).

*In silico* intrinsic terminator prediction was conducted for *B. subtilis* via TransTermHP (*Kingsford et al., 2007*). To ascertain the magnitude of overlap between *B. subtilis* terminators identified in this study, our previous study (*Mondal et al., 2016*), and by TransTermHP (*Kingsford et al., 2007*), we compared the strand identity and 3’ end location of all identified terminators in each population. We considered a terminator to be matched between two Term-seq populations in cases where the strand identity was identical and the 3’ end position of the terminator was within a 4 nt window in both datasets, and matched between a Term-seq population and a TransTermHP population when the strand identity was identical and the 3’ end position of the terminator was within a 15 nt window in both datasets.

### Differential expression analysis and replicate reproducibility

All RNA-Seq reads post-trimming were pseudo-mapped using Kallisto in SE mode using the --rf-stranded option to a transcriptome built with Illumina generated RNA-seq data collected from strain PLBS338 (Supplementary Table 4) (*Bray et al., 2016; Ritchey et al., 2020*). This method determined both TPM and raw count values for each annotated transcript for both merged and non-merged FASTQ files (Supplementary Table 4). TPM values for all coding sequence-containing transcripts were compared for each pair of replicates via both a scatter plot and a spearman correlation analysis (Supplementary Figure 11A). All replicates were found to be highly reproducible with a mean spearman r-value of 0.964. Genes that were differentially expressed between the WT and each mutant strain were identified by analyzing the raw count data from each strain via DESeq2. (Figures 5A, 5B, and 5C) (*Love et al., 2014*) A variance stabilizing transformation was applied to the transcriptome-wide raw count data for each strain, and this matrix was projected onto 2-D space via a principal component analysis (PCA). This analysis revealed that transcriptome-wide expression data collected from each sample clustered neatly by strain (Supplementary Figure 11B).

### Terminator screening and characterization

A 3’ end can be the result of intrinsic termination, Rho-dependent termination, or RNA decay (*Roberts, 2019*). An intrinsic terminator contains a GC-rich RNA hairpin and a U-rich tract immediately downstream of the hairpin (*Roberts, 2019*). Some intrinsic terminators also contain an A-rich tract upstream of the hairpin (*Roberts, 2019*). As such, the 50 nts upstream of each 3’ end was iteratively sent through an *in silico* RNA secondary structure prediction algorithm (RNAStructure) (*Mathews et al., 2004*). In cases where a hairpin was identified, the presence of a U tract leading to the 3’ end was visually queried. Formation of the final two base pairs of a terminator hairpin require the greatest energetic expenditure and are the rate limiting steps of terminator hairpin completion (*Ray-Soni et al., 2016*). As such, bacterial systems have evolved a heavy GC preference at these positions (*Ray-Soni et al., 2016; Peters et al., 2011*). Once the hairpin has completed, termination occurs 7-9 nt downstream from the terminal hairpin nt (*Mondal et al., 2016; Ray-Soni et al., 2016*). These details were factored into intrinsic terminator prediction by assuming that the U-rich tract started with the first U residue for each identified terminator.

Termination efficiency (%T) of a particular intrinsic terminator can be calculated by comparing the median RNA-seq coverage value of the 10 nt upstream (U) to the median RNA-seq coverage value of the 10 nt downstream (D) of the identified 3’ end using the following equation: %T = [(U-D) / U]*(100) as described previously (*Mondal et al., 2016*). Short window sizes of 10 nt were chosen to limit potential complications arising from transcription initiation downstream of the POT. A 3’ end containing the intrinsic terminator modules was included in this study only in cases where %T ≥ 5 in the WT strain. In all cases where the 3’ end position of a terminator identified here did not exactly correspond to the 3’ end position of a matched terminator identified in our previous Term-seq study, the %T was calculated at the 3’ end position identified previously (*Mondal et al., 2016*).

To determine the effect of an elongation factor, or combination of elongation factors, on the %T of a particular intrinsic terminator, one can calculate the change in termination efficiency (Δ%T) using the following equation: Δ%T = %T_WT_ - %T_mutant_. This approach was systematically applied to all intrinsic terminators to determine the general effect of an elongation factor, or combination of elongation factors, on intrinsic termination (Supplementary Table 2). Determination of all terminator hairpin stem lengths and loop lengths were derived from the result of sending the predicted terminator hairpin stem through the *in silico* RNA secondary structure prediction algorithm (RNAStructure) (*Mathews et al., 2004*).

### DNA templates and plasmids

All pAY196 derivatives were generated using a strategy akin to site-directed mutagenesis PCR (*Hemsley et al., 1989*). The entirety of pAY196 was amplified using Vent polymerase (New England Biolabs) using outward-directed primer pairs with constant sequences complementary to the plasmid backbone and flanking regions containing the biological sequence of interest as described previously (*Mondal et al., 2016*). To prevent the formation of primer dimers or internal hairpins caused by terminator hairpins, the biological sequence of one primer contained an A tract when present and the 5’ portion of the predicted hairpin ending at the 3’ most nt of the loop. The biological sequence of the other primer contained the 3’ portion of the predicted hairpin and 19 nt downstream of the predicted hairpin. All plasmids and primers used in this study are listed in Supplementary Table 7 and Supplementary Table 8, respectively.

### *In vitro* transcription

Analysis of RNAP pausing and termination was performed as described previously with modifications (*Mondal et al., 2017*). DNA templates were PCR-amplified from plasmids containing either WT or mutant terminator sequence, both of which included the predicted terminator hairpin, 19 nt downstream of the predicted hairpin, and the A tract when present, fused to the *B. subtilis* P_*trp*_ promoter and *trp* leader derived C-less cassette (pAY196 and derivatives) using PSL (modifies the P_*trp*_ promoter to a consensus promoter with an extended −10 element) and lacZ primers (Supplementary Figure 12A). Halted elongation complexes containing a 27-nt transcript were formed for 5 min at 37°C by combining equal volumes of 2X template (50-200 nM) with 2X halted elongation complex master mix containing 80 μM ATP and GTP, 2 μM UTP, 100 μg/ml bovine serum albumin, 150 μg/ml (0.38 μM) *B. subtilis* RNAP holoenzyme, 0.76 μM SigA, 2 μCi of [α-^32^P]UTP and 2X transcription buffer (1X = 40 mM Tris-HCl, pH 8.0, 5 mM MgCl_2_, 5% trehalose, 0.1 mM EDTA, and 4 mM dithiothreitol). RNAP and SigA were added from a 20x stock solution containing 1.5 mg/ml RNAP and 0.35 mg/ml SigA in enzyme dilution buffer (1X = 20 mM Tris-HCl, pH 8.0, 40 mM KCl, 1 mM dithiothreitol, and 50% glycerol). A 4X solution containing either 0 μM NusA and 0 μM NusG, 4 μM NusA and 0 μM NusG, 0 μM NusA and 4 μM NusG, or 4 μM NusA and 4 μM NusG in 1X transcription buffer were added, and the resulting solution was incubated for 5 min at 23°C. For termination assays, a 4X extension master mix containing 80 μM KCl, 600 μM of each NTP, 400 μM rifampicin, in 1X transcription buffer was added, and the reaction was allowed to proceed for 30 min at 23°C before the addition of an equal volume of 2X stop/gel loading solution (40 mM Tris-base, 20 mM Na_2_EDTA, 0.2% sodium dodecyl sulfate, 0.05% bromophenol blue, and 0.05% xylene cyanol in formamide). For pausing assays, the same extension master mix was added, and the reaction was incubated at 23°C, with aliquots removed and stop/gel loading solution added at the specified time points. A 30 min time point was included for all pausing assays that mirrored the experimental conditions of the termination assay. RNA bands were separated on standard 5% sequencing polyacrylamide gels. All RNA sequencing reactions were conducted like other termination reactions, albeit with the addition of one of four 3’ dNTPs at a 1:1 molar ratio with the corresponding NTP within the extension master mix. Termination efficiencies and pausing half-lives were quantified as described previously (*Yakhnin et al., 2002*). Each *in vitro* experiment was conducted a minimum of 2 times, with representative gels and termination efficiencies/pausing half-lives shown in the figures.

### Convergent transcription *in vitro*

gBlock gene fragments (IDT) containing biological sequence (100 nt upstream of, 50 nt downstream of the predicted *serA* and *fisB* terminator hairpins) flanked by two identical 27-nt C-less cassettes, inward facing consensus promoters with extended −10 elements (P_forward_ and P_reverse_), and EcoRI and HindII restriction digestion sites (NEB) were cloned into pTZ19R (Thermo Fisher). DNA templates containing both P_forward_ and P_reverse_ were PCR-amplified from the appropriate pTZ19R derivative using a primer pair that was specific for pTZ19R (M13_2.0 and M13_reverse_2.0, IDT, derivatives of M13 and M13 Reverse universal sequencing primers). Conversely, DNA templates containing just P_forward_ were PCR-amplified from the appropriate pTZ19R derivative using a reverse primer specific for pTZ19R (M13_2.0) and a forward primer specific for the biological sequence (serA_uni/fisB_uni, Supplementary Figure 12B). *In vitro* transcription termination reactions were conducted identically using both of the templates detailed in Supplementary Figure 12.

### Motility assay

Swimming motility assays were conducted as performed previously with modifications (*Mukherjee et al., 2016*). Strains were grown to mid-exponential phase in the presence of 0.2 mM IPTG and concentrated to 10 OD_600_ in phosphate-buffered saline (PBS) pH (137 mM NaCl, 2.7 mM KCl, 10 mM Na_2_HPO_4_, and 2 mM KH_2_PO_4_). LB plates containing 0.3% Bacto agar ± 0.2 mM IPTG were dried for 10 min in a laminar flow hood, centrally inoculated with 10 μL of the cell suspension, dried for another 10 min, and incubated for ~13 hr at 37°C in a humid chamber. Plates were visualized with a BioRad Geldoc system and digitally captured using BioRad Quantity One software.

### Microscopy

Fluorescence microscopy was conducted with a Nikon 80i microscope with a phase contract objective Nikon Plan Apo 100X and an Excite 120 metal halide lamp as described previously with modifications (*Mukherjee et al., 2016*). FM4-64 (Molecular Probes) was visualized with a C-FL HYQ Texas Red Filter Cube (excitation filter 532-587 nm, barrier filter > 590 nm). Alexa 488 C_5_ maleimide (Molecular Probes) fluorescent signals were visualized using a C-FL HYQ FITC Filter Cube (FITC, excitation filter 460 – 500 nm, barrier filter 515-550 nm).

To visualize flagella, cells were grown at 37°C in LB broth + 0.2 mM IPTG to mid-exponential phase. One mL of broth culture was harvested and resuspended in 50 μL of PBS containing 5 μg/mL Alexa Fluor 488 C_5_ maleimide (Molecular Probes), and incubated for 3 min at 23°C as described previously (*Blair et al., 2008*). Cells were then washed with 1 mL of PBS.

Membranes were stained by resuspension of 30 μL of PBS containing 5 μg/mL FM4-64 and incubated for 5 min at 23°C, then washed with 1 mL of PBS. Cell pellets were resuspended with 30-50 μL of PBS, and then 4 μL of suspension were placed on a microscope slide and immobilized with a poly-L-lysine treated coverslip. For cells to be depleted of NusA (PLBS730 and PLBS731), cells were initially grown in LB supplemented with 0.2 mM IPTG to mid-exponential phase. One mL of culture was harvested, washed 2X with fresh LB, back diluted in fresh LB and grown for 4 generations at 37°C. One mL of culture was harvested and staining of the flagella and membrane was conducted via the above protocol.

## Data Availability

RNA-seq libraries were deposited in Gene Expression Omnibus (GEO) (accession number GSE154522). The reviewer access code is urersmoqjluflwh. All other data analyzed during this study are included in this published article (and its supplementary information files).

## Code Availability

All custom scripts used for 3’ end mapping are available at https://github.com/zfmandell/Term-seq and all other scripts are available upon request.

The authors declare no competing interests.

## Acknowledgements

Illumina sequencing was performed at the Penn State Genomics Core Facility. This work was supported by National Institutes of Health Grant GM098399 to Paul Babitzke, National Institutes of Health Grant GM131783 to Daniel B. Kearns, and the Intramural Research Program of the National Institutes of Health/National Cancer Institute to Mikhail Kashlev.

## Author Contributions

Z.F. Mandell performed Term-seq, RNA-seq, bioinformatics analyses, western blotting, *in* vitro transcription, interpreted data, prepared figures, and wrote the manuscript. R.T. Oshiro performed motility assays and microscopy. A.V. Yakhnin contributed to experimental design and provided guidance for *in vitro* transcription. M. Kashlev interpreted data. D.B. Kearns contributed to experimental design and directed the motility and microscopy components of the project. P. Babitzke conceived and directed the project, wrote the manuscript, analyzed data and prepared figures.

---

**Figure S1.**
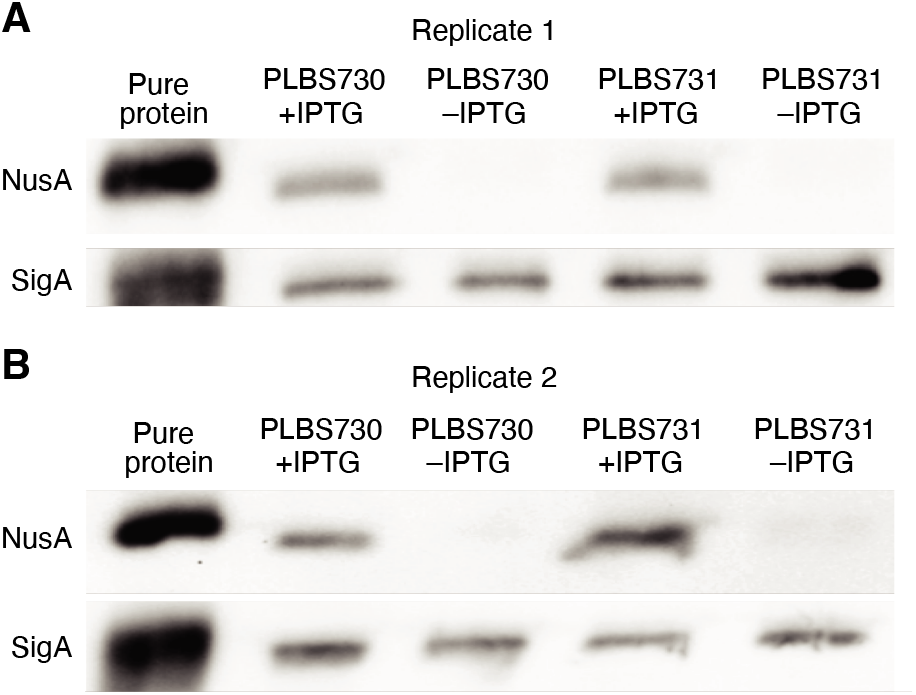
Western blot analysis of NusA depletion. (**A**) Western blot for all samples used for Term-seq replicate 1. Top panel, image after probing for NusA. Bottom panel, image after probing for SigA as a loading control. Purified NusA and SigA are in the left lane. Lanes containing protein extracted from strains PLBS730 ± IPTG and PLBS731 ± IPTG are specified. (**B**) Identical to panel **A** except samples from replicate 2.

**Figure S2.**
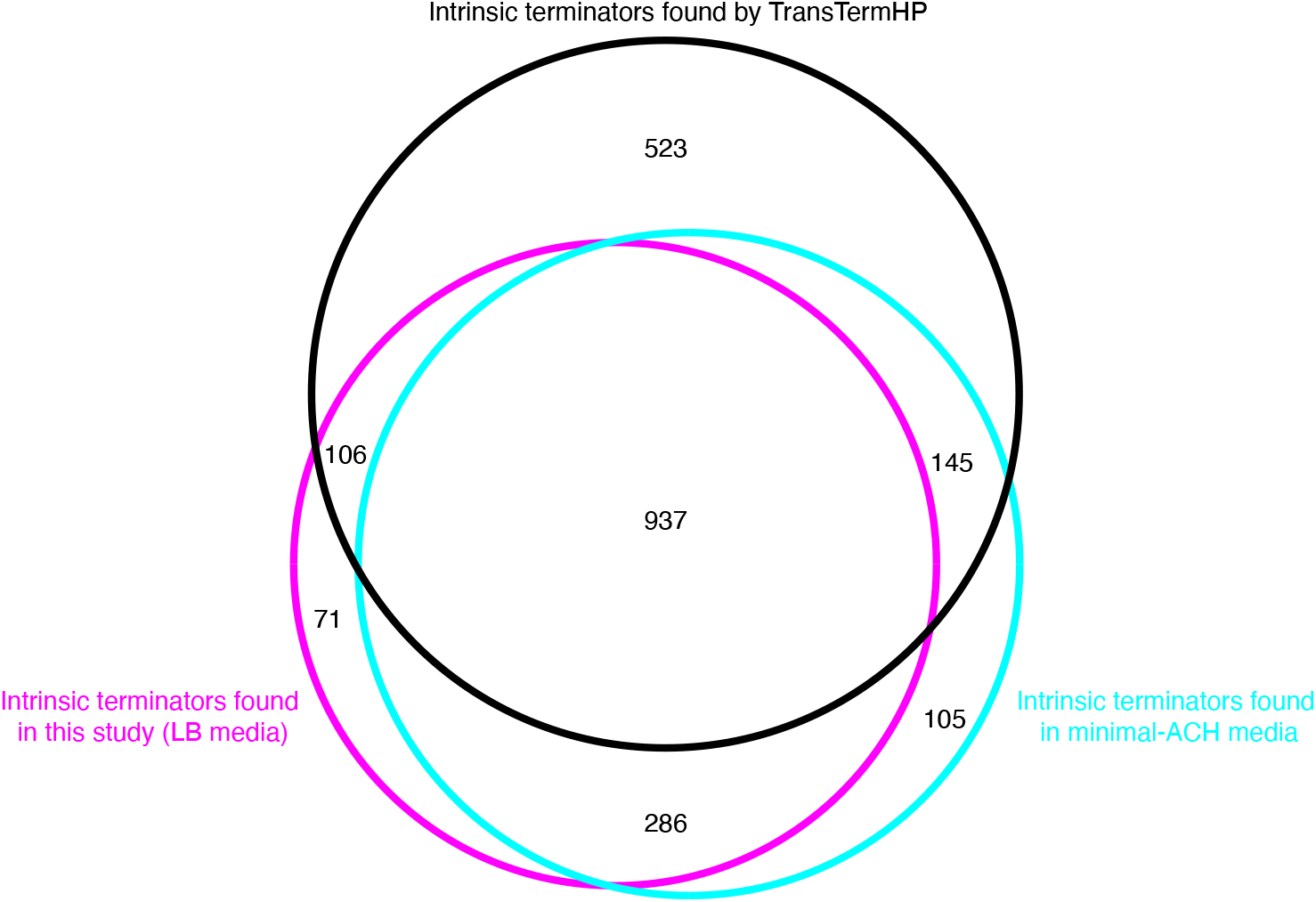
Benchmarking the intrinsic terminators identified in this study. Venn diagram showing the number and overlap of *B. subtilis* terminators identified within three distinct datasets. The black circle contains all terminators that were identified by the *in silico* tool TransTermHP, the magenta circle contains all terminators that were identified here via Term-seq conducted in WT *B. subtilis* grown in LB media, and the cyan circle contains all terminators that were identified previously via Term-seq conducted in WT *B. subtilis* grown in Minimal-ACH media (*Mondal et al., 2016*).

**Figure S3.**
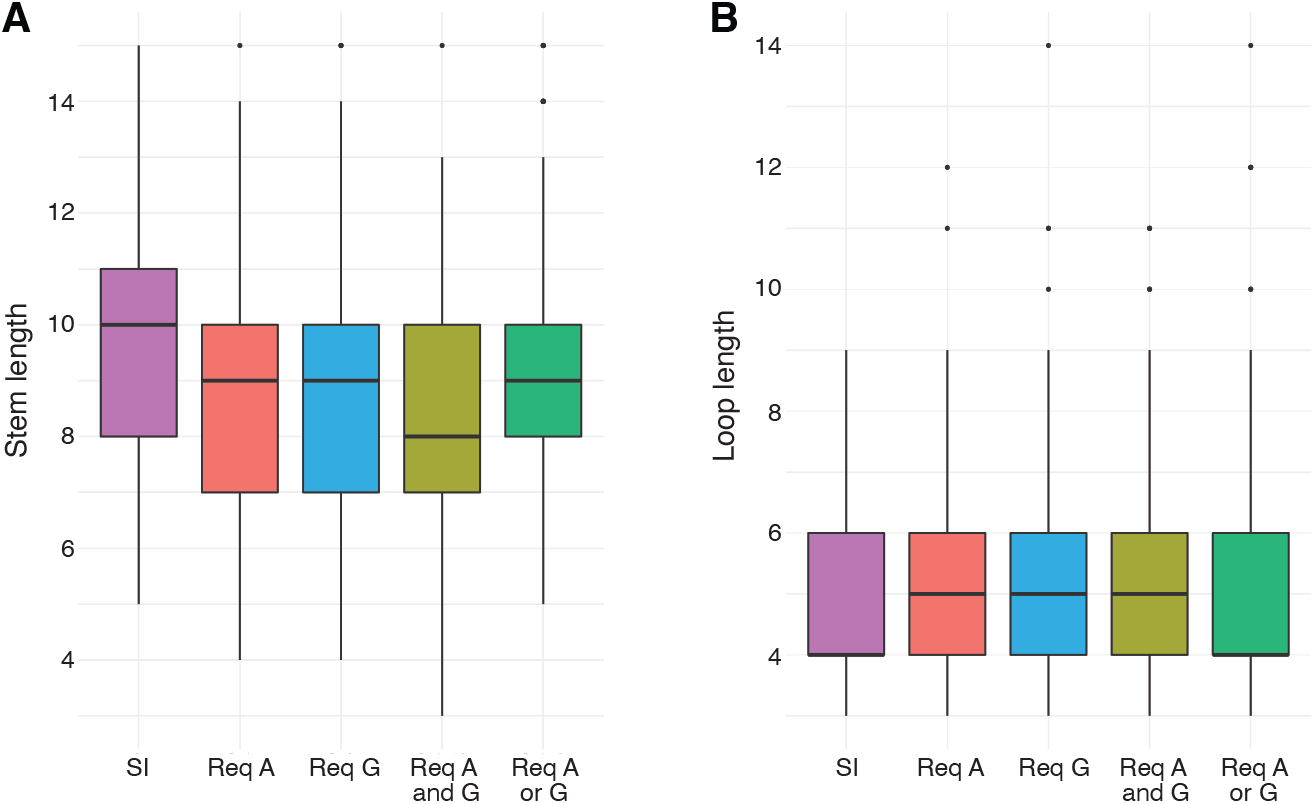
Intrinsic terminator hairpin stem length and loop length. (**A**) Box plot showing the distribution of termination hairpin stem lengths of the SI, Req A, Req G, Req A and G, and Req A or G terminator subpopulations identified in Figure 1B. (**B**) Box plot showing the distribution of termination hairpin loop lengths of the SI, Req A, Req G, Req A and G, and Req A or G terminator subpopulations identified in Figure 1B.

**Figure S4.**
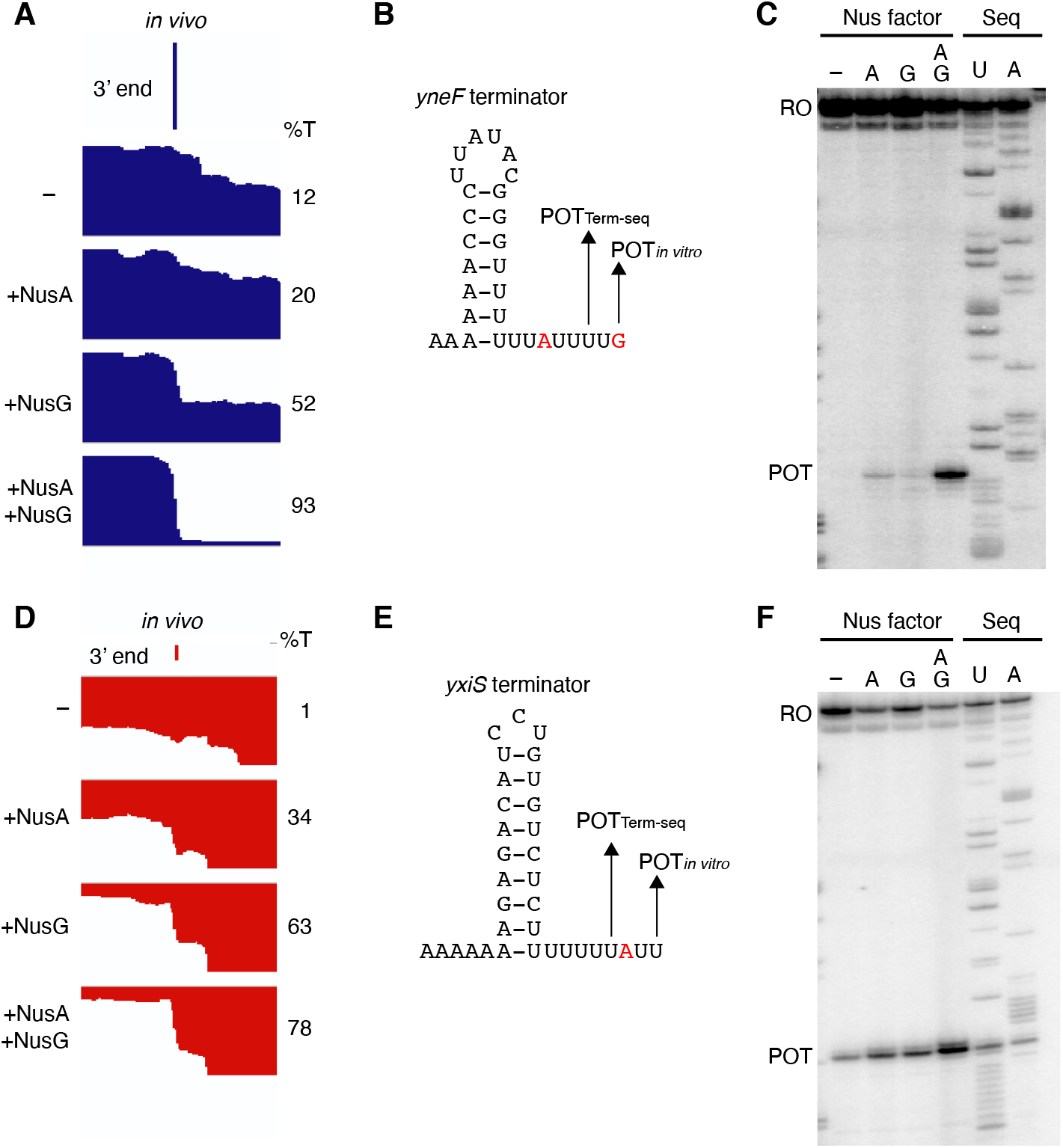
NusG stimulates termination at terminators containing A-U base pairs at the base of the hairpin. (**A**) IGV screenshot of a genomic window centered around the *yneF* terminator. Top track is the 3’ end identified by Term-seq. Bottom tracks are the RNA-seq coverage data for the *nusA*dep Δ*nusG* (−), Δ*nusG* (+NusA), *nusA*dep (+NusG), and WT (+NusA +NusG) strains. %T in each strain is shown on the right of each track. Transcription proceeds from left to right. (**B**) *yneF* terminator showing the point of termination identified *in vivo* by Term-seq (POT_Term-seq_) and by *in vitro* transcription in the +A+G condition (POT_*in vitro*_). Disruptions in the U-rich tract are shown in red. The upstream A tract is also shown. (**C**) Single-round *in vitro* termination assay with the *yneF* terminator. Experiments were performed in the absence (−) or presence of NusA (A) and/or NusG (G) as indicated. Positions of terminated (POT) and run-off (RO) transcripts are marked. RNA sequencing lanes (U, A) are labeled. (**D-F**) Identical to panels **A-C** except that it is for the *yxiS* terminator. (**D**) Transcription is from right to left.

**Figure S5.**
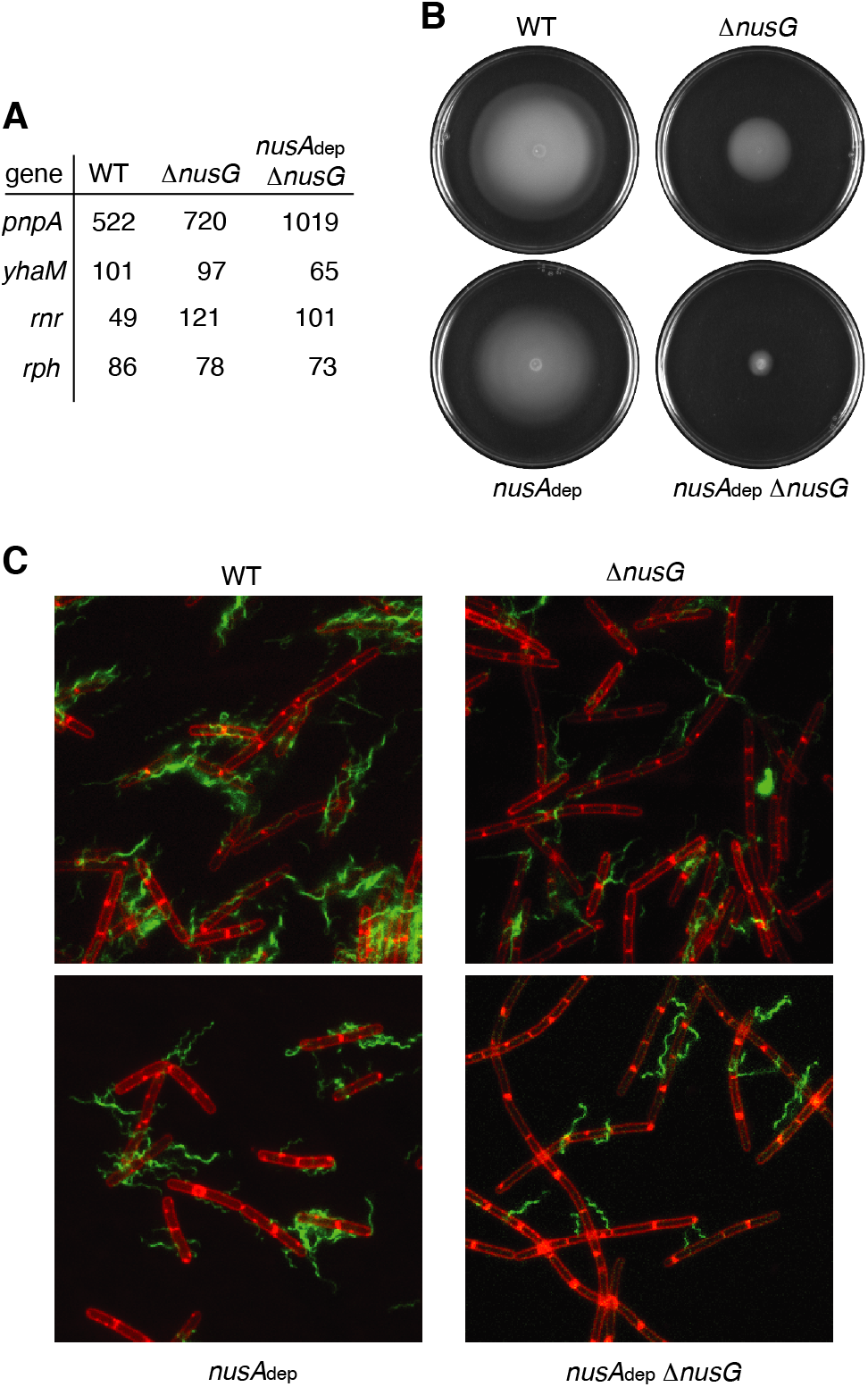
NusG is a motility factor in *B. subtilis.* (**A**) TPM values calculated for *pnpA*, *yhaM, rnr,* and *rph*, four genes that encode 3’ to 5’ exoribonuclease, in WT, Δ*nusG*, and *nusA*dep Δ*nusG* strains. (**B**) Swimming motility assay for WT, *nusA*dep, Δ*nusG*, and *nusA*dep Δ*nusG* strains. (**C**) Florescence microscopy performed on WT, *nusA*dep, Δ*nusG*, and *nusA*dep Δ*nusG* strains. Membrane is stained with FM4-64 (false colored red) and flagella are stained with Alexa Fluor 488 C5 maleimide (false colored green).

**Figure S6.**
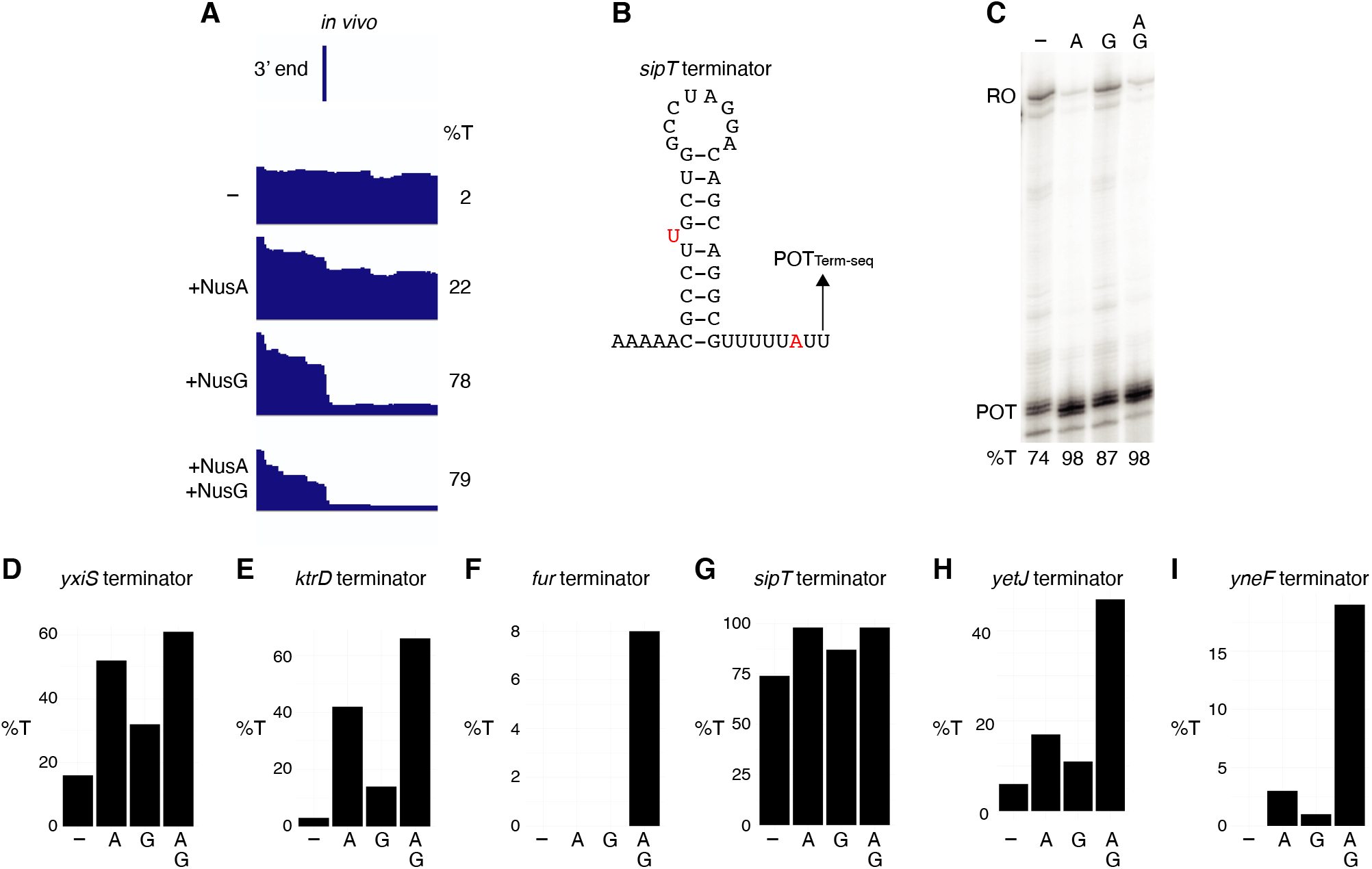
NusA is a more potent termination factor *in vitro* than NusG. (**A**) IGV screenshot of a genomic window centered around the *sipT* terminator. Top track is the 3’ end identified by Term-seq. Bottom tracks are the RNA-seq coverage data for the *nusA*dep Δ*nusG* (−), Δ*nusG* (+NusA), *nusA*dep (+NusG), and WT (+NusA +NusG) strains. %T in each strain is shown on the right of each track. Transcription proceeds from left to right. (**B**) *sipT* terminator showing the point of termination identified *in vivo* by Term-seq (POT_Term-seq_). Disruptions in the hairpin and U-rich tract are shown in red. The upstream A tract is also shown. (**C**) Single-round *in vitro* termination assay with the *sipT* terminator. Experiments were performed in the absence (−) or presence of NusA (A) and/or NusG (G) as indicated. Positions of terminated (POT) and run-off (RO) transcripts are marked. %T is shown below each lane. (**D-I**) *In vitro* termination efficiencies for the specified terminators in the absence (−) or presence of NusA (A) and/or NusG (G) as indicated.

**Figure S7.**
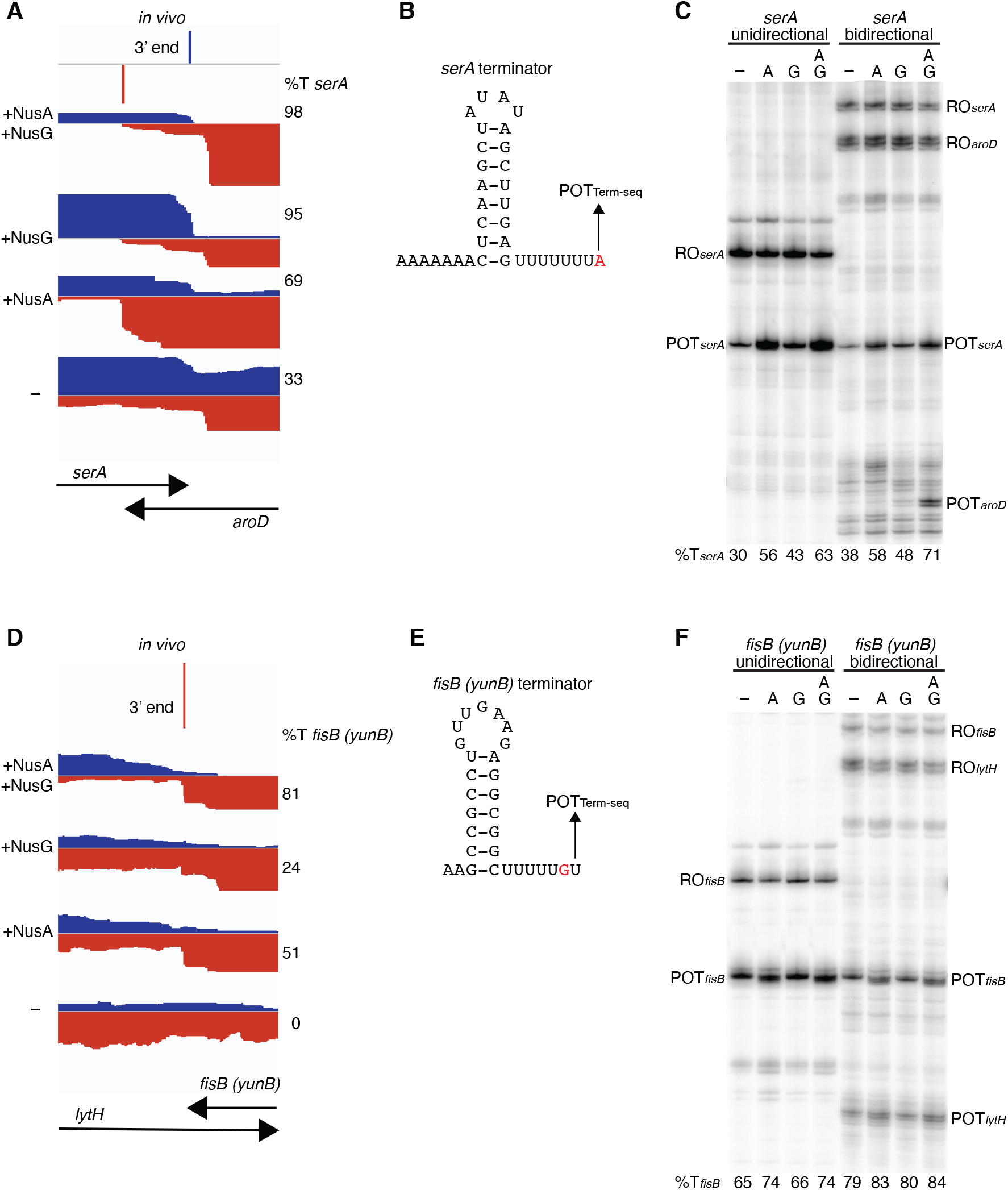
Convergent transcription does not modify the impact of NusG in vitro. (**A**) IGV screenshot of a genomic window showing the intersection of *serA* and *aroD*. Top track is the 3’ end of *serA* identified by Term-seq. Bottom tracks are the RNA-seq coverage data for the WT (+NusA +NusG), *nusA*dep (+NusG), Δ*nusG* (+NusA), and *nusA*dep Δ*nusG* (−) strains. %T in each strain is shown on the right of each track. Transcription proceeds from left to right for *serA* (blue) and right to left for *aroD* (red). (**B**) *serA* terminator showing the point of termination identified *in vivo* by Term-seq (POT_Term-seq_). The upstream A tract is also shown. (**C**) *In vitro* transcription of the *serA* terminator (*serA* unidirectional) or both the *serA* and *aroD* terminators (*serA* bidirectional). Experiments were performed in the absence (−) or presence of NusA (A) and/or NusG (G) as indicated. Positions of terminated (POT) and runoff (RO) products are marked. %T is shown below each lane. (**D-F**) Identical to panels **A-C** except that it is the *fisB* (*yunB*) terminator at the *fisB-lytH* intersection.

**Figure S8.**
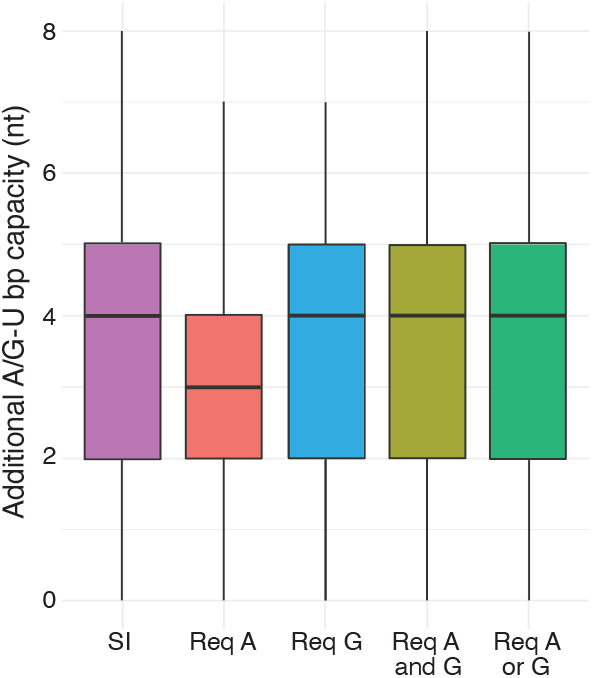
Capacity for additional A/G-U base pairing in terminator subpopulations. Box plot showing the distribution of the capacity for additional A/G-U base pairing of the SI, Req A, Req G, Req A and G, and Req A or G terminator subpopulations identified in Figure 1B.

**Figure S9.**
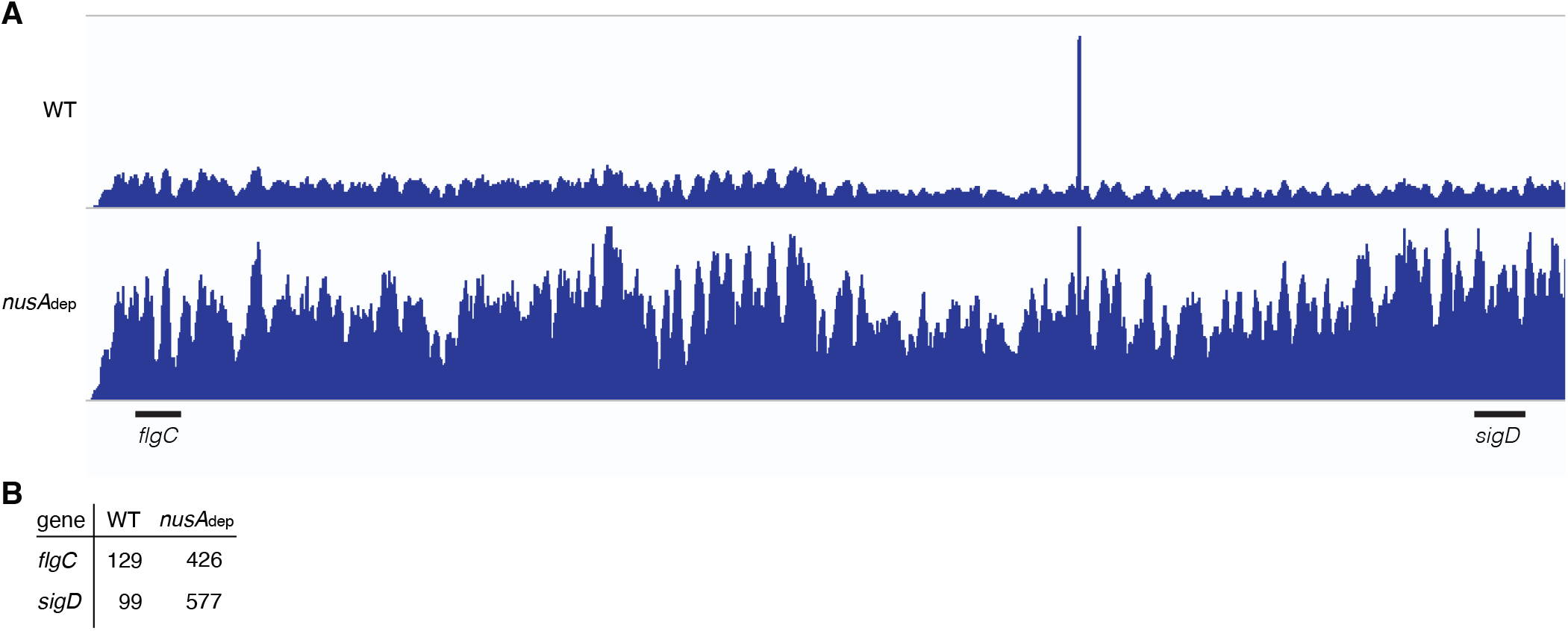
NusA may serve as a transcription destabilization factor. (**A**) IGV screenshot of the fla/che operon. Each track is the RNA-seq coverage data for the WT and *nusA*dep strains. Locations of the *flgC* and *sigD* genes are specified below the screenshot. (**B**) TPM values calculated for the *flgC* and *sigD* genes in the WT and *nusA*dep strains.

**Figure S10.**
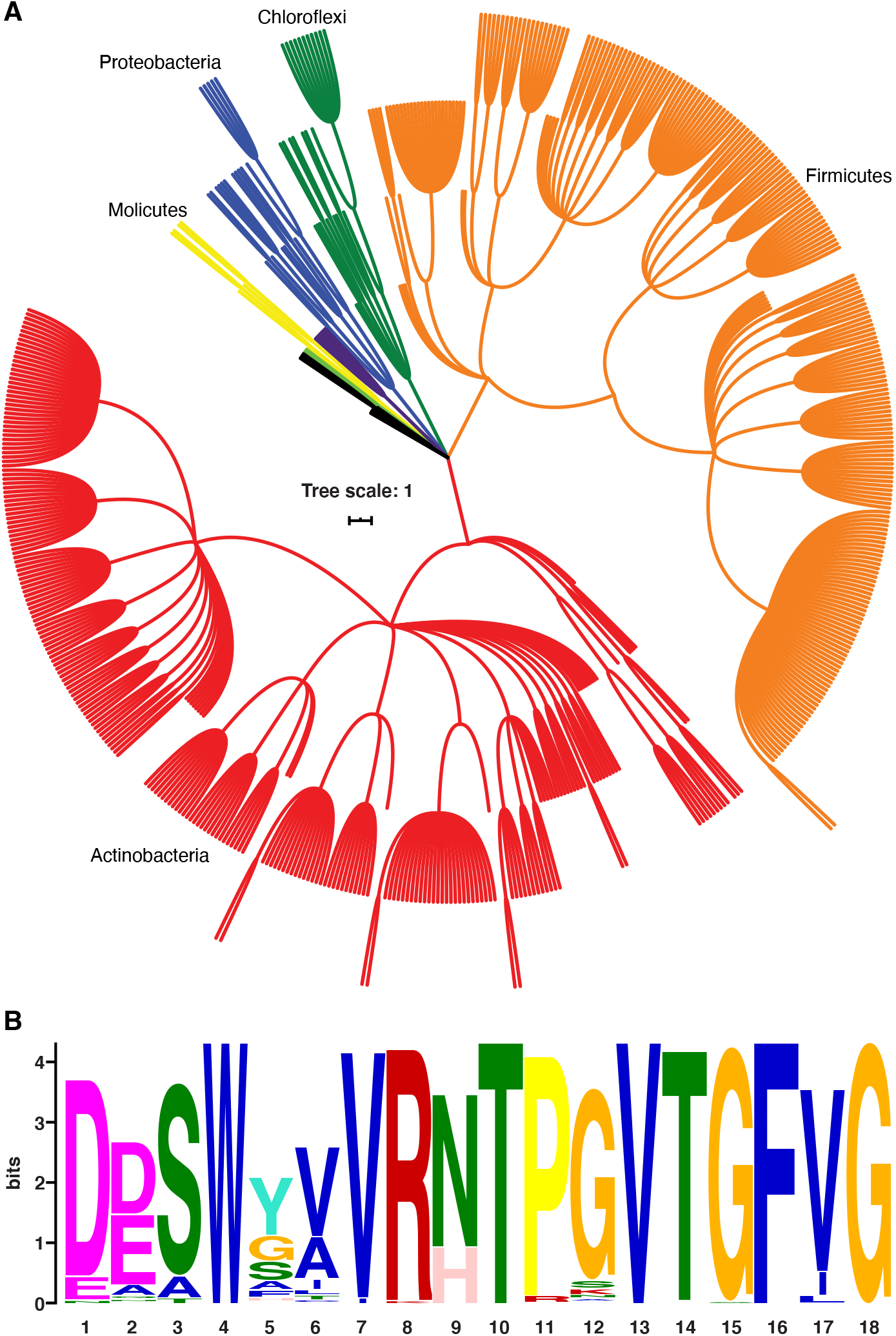
NusG homologs exhibit the capacity to contact the ntDNA strand across the bacterial domain. (**A**) Phylogenetic tree constructed from the 16S rRNA sequences of all bacteria found to encode a NusG homolog that contains the *B. subtilis*-like dipeptide residues (NT and HT). Each different phylum of bacteria that is represented with 3 or more species is highlighted a different color. Actinobacteria branches are red, Firmicute branches are orange, Chloroflexi branches are dark green, Proteobacteria branches are blue, Synergistaceae branches are purple, Mollicute branches are yellow, and Bacteroidete branches are light green. Phyla with fewer than three representative species are in black. (**B**) Sequence logo constructed from the portion of NusG that interacts with the ntDNA strand in *B. subtilis* for all NusG homologs present in **A.** Critical dipeptide NT/HT is located at positions 9 and 10.

**Figure S11.**
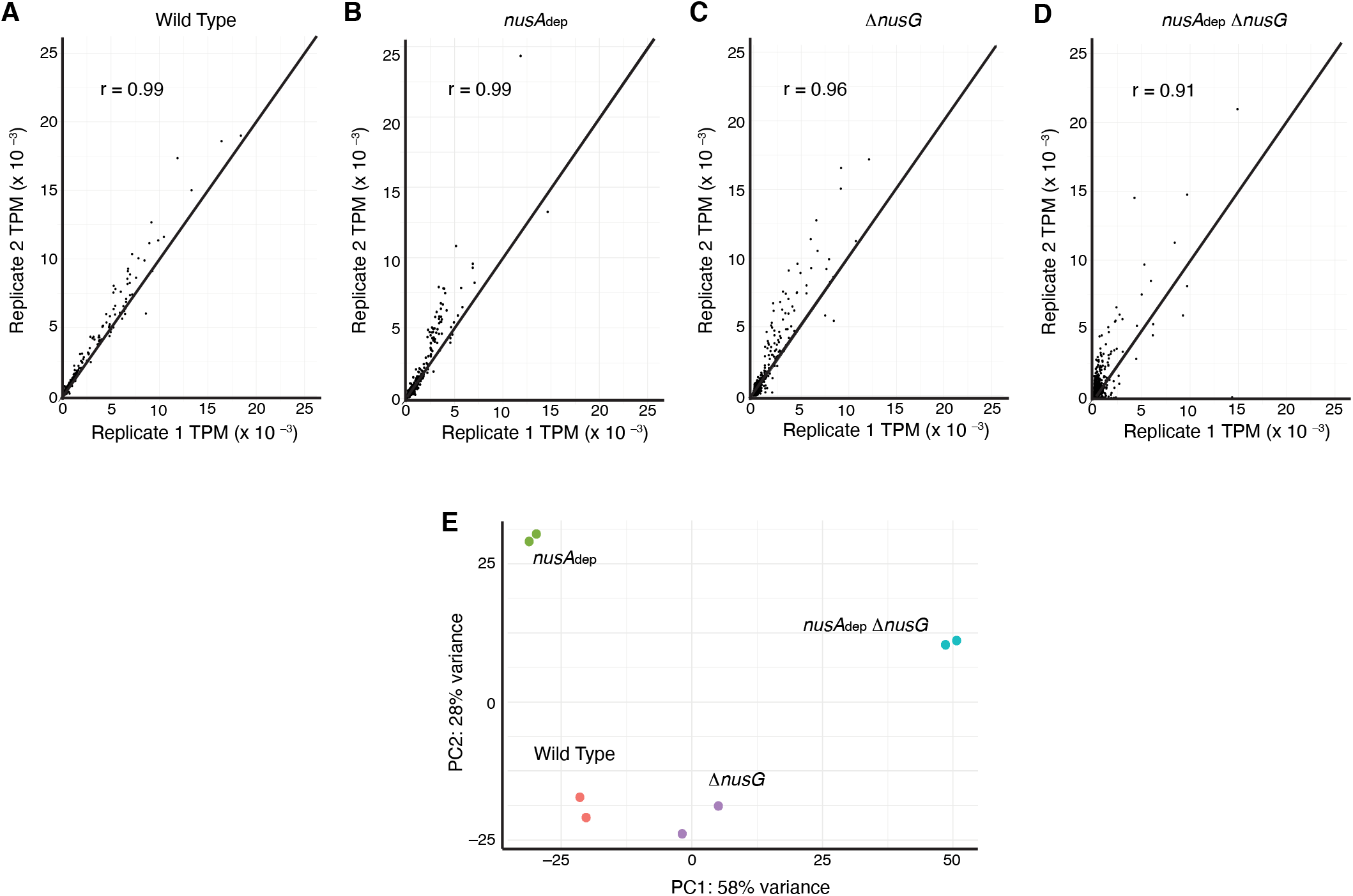
Transcriptomics data showing that Term-seq replicates are highly correlated and data from each strain is distinct. (**A**) Scatter plot of all protein coding sequence TPM values calculated from the WT strain replicates. Line is through the diagonal (y = x) and spearmen correlation value specified in the plot. (**B**) Identical to panel **A** except for *nusA*dep strain replicates. (**C**) Identical to panel **A** except for Δ*nusG* strain replicates. (**D**) Identical to panel **A** except for *nusA*dep Δ*nusG* strain replicates. (**E**) Principal component analysis (PCA) plot of transcriptomics data collected from each sample.

**Figure S12.**
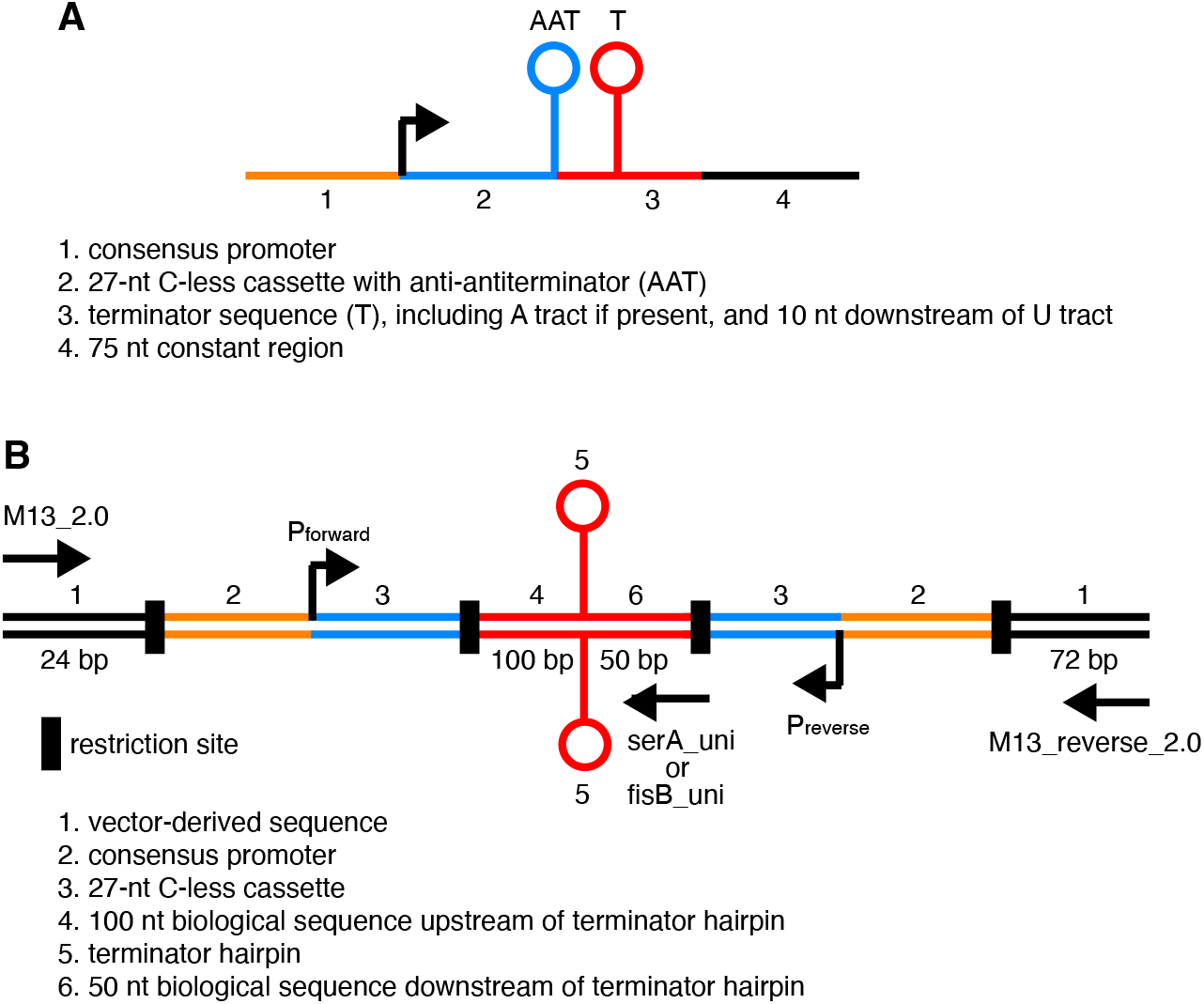
Template design for in vitro transcription. (**A**) Schematic representation of templates used for all *in vitro* transcription experiments except for those involving convergent transcription. Template features are described below. (**B**) Schematic representation of templates used for *in vitro* transcription experiments that involve convergent transcription. Template features are described below including locations of restriction sites and primer annealing for the generation of templates.

